# Unraveling the Impact of W215A/E217A Mutations on Thrombin’s Dynamics and Thrombomodulin Binding through Molecular Dynamics Simulations

**DOI:** 10.1101/2023.12.20.572552

**Authors:** Dizhou Wu, Freddie R. Salsbury

## Abstract

Thrombin, a central serine protease in hemostasis, exhibits dual functionality in coagulation processes—favoring fibrinogen cleavage in its native form while shifting towards protein C activation when complexed with thrombomodulin (TM). Thrombin also plays roles in cancer-associated thrombosis and may be involved in metastasis and tumorigenesis. The W215A/E217A (WE) double mutant of thrombin presents a unique case, with its fibrinogen cleavage activity diminished by 19,000-fold, contrasting a modest 7-fold reduction in protein C activation in the presence of TM. The differential substrate specificity of this mutant raises fundamental questions about the underlying molecular mechanisms. In this study, we employed all-atom microsecond-scale molecular dynamics (MD) simulations, complemented by Root Mean Square Fluctuation (RMSF) analysis, clustering algorithms, PCA-based free-energy surfaces, and logistic regression modeling, to dissect the structural and allosteric changes driving thrombin’s substrate specificity. Our results unveil distinct conformational states within the catalytic triad, each optimized for specific substrate interactions. We demonstrate that the WE mutations synergize with TM456 binding, resulting in altered hydrogen bond networks and distinct free energy landscapes. A key finding of our research is the identification of ARG125 as a pivotal element in these interactions, consistently forming critical hydrogen bonds across different thrombin variants. The persistent role of ARG125 not only elucidates aspects of thrombin’s functional plasticity but also positions it as a promising target for novel therapies. This comprehensive analysis enhances our understanding of thrombin’s structural dynamics, paving the way for more effective and targeted therapeutics.

## Introduction

Thrombin, a key serine protease in the coagulation cascade, plays a dual role in hemostasis. It is essential for converting fibrinogen into fibrin, thus facilitating clot formation[1, 2]. Moreover, thrombin engages in anticoagulant pathways, particularly in its interaction with protein C, an interaction enhanced by the endothelial cell cofactor thrombomodulin (TM)[1, 2]. Thrombin’s binding to TM is a pivotal event, altering its substrate specificity from a procoagulant agent—by inhibiting fibrinogen binding—to promoting anticoagulant activity through protein C activation[1, 2]. This modulation of thrombin’s function is crucial in maintaining the balance between procoagulant and anticoagulant mechanisms, vital for hemostatic equilibrium[1, 2]. Thrombin is also associated with thrombosis in cancer[3] and may play a role in metastasis and tumorigenesis [4]. Significant interest has been garnered by the W215A/E217A (WE) double mutant of thrombin, which exhibits profound anticoagulant properties. This mutant, involving modifications at chymotrypsin numbering residues 215 and 217 (equivalent to 263 and 265 in sequential numbering), has been studied extensively for its altered coagulation profile[5, 6, 7, 8, 9, 10, 11]. The WE variant marks a significant shift in thrombin’s function, favoring anticoagulant activities over procoagulant ones[12, 13]. Notably, it demonstrates an over 19,000-fold reduction in fibrinogen cleavage capacity, compared to a 7-fold decrease in protein C activation in conjunction with TM[14, 5, 15, 16]. This paper adheres to sequential numbering for residue identification.

Crystallographic studies have shown that the WE mutation collapses thrombin’s primary substrate binding pocket[17], significantly impairing its procoagulant activity. Despite this, the WE mutant maintains its ability to cleave protein C when bound to TM, an intriguing phenomenon given the compromised binding pocket. Recent insights from hydrogen/deuterium exchange mass spectrometry (HDXMS) indicate that TM may reshape the active sites of the WE mutant, orienting it towards protein C activity[16]. Yet, the detailed atomic-level dynamics of TM’s modulation of WE thrombin are not fully understood.

To elucidate the functional retention in the WE mutant, our study employs molecular dynamics (MD) simulations, supplemented by analytical techniques such as Root Mean Square Fluctuation (RMSF), clustering algorithms, Principal Component Analysis (PCA), and logistic regression models. We aim to explore the structural dynamics and allosteric regulation in various thrombin variants, including wild-type, TM456-bound wild-type, WE, and TM456-bound WE.

Our analysis reveals distinct conformational states within the catalytic triad, each optimized for specific substrate interactions, as indicated by HDBSCAN clustering and supported by hydrogen bonding analyses[18, 15, 17, 5]. The WE mutations appear to cause an allosteric shift in the catalytic triad towards protein C activation, possibly explaining the significant reduction in fibrinogen cleavage. In TM’s presence, the WE mutant maintains this functional bias but exhibits unique structural features in the catalytic triad and other regions, leading to only a minor decrease in protein C activation.

Additionally, our findings highlight a synergistic effect between the WE mutations and TM456 binding, characterized by novel hydrogen bonding patterns and altered conformational landscapes. Particularly, ARG125 emerges as a key hydrogen bond participant across all thrombin variants, underscoring its potential as a therapeutic target.

## Materials and methods

### Simulation System

To explore the catalytic efficiency of the WE thrombin variant towards protein C in the presence of TM, despite its compromised binding pocket, we conducted molecular dynamics (MD) simulations under four different scenarios: wild-type thrombin, TM456-bound wild-type thrombin, the WE double mutant thrombin, and TM456-bound WE double mutant thrombin. These simulations aimed to develop a comprehensive conformational ensemble of thrombin, delineating the effects of both TM456 binding and mutational alterations. Through comparative analysis, we intend to pinpoint crucial conformational changes and vital hydrogen bonds that facilitate the sustained catalytic function of the WE thrombin variant with TM.

Our study employed high-resolution crystal structures of the TM456-thrombin complex, available from the RCSB Protein Data Bank (PDB ID: 1DX5), as resolved by Fuentes-Prior et al. in 2000[19]. This structure has been the subject of several other computational studies. [20, 21, 22, 23, 24, 25, 26, 27] The original structure, however, included several non-physiological mutations: ARG456 and HIS457 were altered to GLY456 and GLN457, respectively, to inhibit proteolysis[19]; ASN364 was converted to ASP364 post PNGase treatment[19]; and THR91 was modified to ILE91. To align our simulations with physiological conditions, we reverted these residues to their native states in the wild-type and TM456-bound wild-type simulations. In contrast, for the WE and TM456-bound WE simulations, TRP263 and GLU265 were mutated to ALA. The initial PDB file was refined to include only the essential components - the light chain, heavy chain, TM456, and calciumion - discarding all other ligands and water molecules. Missing hydrogen atoms were added utilizing the psfgen tool in VMD, following standard parameters[28].

### Simulation Conformations

To encompass a wide array of conformational states, we conducted eight independent 1-µs all-atom molecular dynamics (MD) simulations for each thrombin system variant: wild-type, TM456-bound wild-type, double mutant, and TM456-bound double mutant. These simulations were performed using the ACEMD3 simulation package, renowned for its GPU acceleration capabilities, and were run on NVIDIA Titan GPUs housed in Metrocubo workstations[29]. Our simulation protocols are consistent with ACEMD’s standards and build on the methodologies validated in our previous thrombin-centric and other biological studies[20, 21, 22, 23, 24, 25, 26, 27, 30, 31, 32, 33, 34, 35, 36, 37].

Each thrombin system was immersed in an explicit TIP3P cubic water box, ensuring a minimum buffer of 10 Å in all directions[38]. Electrical neutrality was achieved by adding Cl ions. The ionic strength was then adjusted to a NaCl concentration of 0.125 M, adhering to our established simulation protocols[24, 25, 26, 27, 20, 21, 22, 23]. All ionizable residues were set to their default protonation states, reflecting a physiological pH of 7. To mitigate any potential steric clashes, each system was subjected to 1000 cycles of conjugate gradient minimization prior to the simulation runs.

The CHARMM36 force field was employed for these simulations[39]. We maintained a constant pressure of 1 atm using the Berendsen pressure control method[40] and regulated the temperature at 300 K using the Langevin thermostat, set with a damping coefficient of 0.1[41]. Van der Waals and electrostatic interactions were calculated up to a cutoff of 9 Å, with a switching distance set at 7.5 Å. Long-range electrostatics were handled using the smooth-particle-mesh-Ewald (PME) method[42, 43].

A 4-fs time step was utilized for numerical integration, enabled by hydrogen mass repartitioning[44]. The SHAKE algorithm constrained bond lengths, allowing for this extended integration step[45]. We recorded system conformations at every 10-ps interval, resulting in 100,000 frames for each 1-*µ*s simulation.

In total, this methodology yielded 800,000 frames per system across the eight 1-µs simulations. These 32 trajectories were then sequentially concatenated, respecting the order of wild-type, TM456-bound wild-type, double mutant, and TM456-bound double mutant, forming a cumulative 32-*µ*s trajectory of 32,000 frames. For analysis, all frames were aligned to the initial thrombin structure through rigid body transformations, using a custom Python script[46]. This alignment facilitated a focused analysis of internal thrombin dynamics across all subsequent examinations.

### Root Mean Square Fluctuations

In our analysis, we employed Root Mean Square Fluctuations (RMSFs) to quantitatively evaluate atomic flexibility throughout the simulations. RMSFs are a crucial metric for determining the average spatial deviation of individual atoms from their mean positions, offering a window into the dynamic properties and relative flexibility of specific protein regions. For clarity, it should be noted that, unless otherwise specified, the RMSFs discussed are for the alpha carbons of the residues. The RMSF for a given atom *i* is calculated according to the following equation:

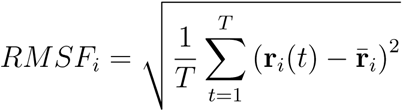

Here, **r***_i_*(*t*) represents the position vector of atom *i* at frame *t*, and **r̄***_i_* denotes the time-averaged position of atom *i* across all *T* frames.

### Clustering Analysis

To decipher the conformational dynamics of thrombin under the influence of TM456 binding and double mutations, we employed non-parametric clustering analysis. This technique is fundamental for systematically categorizing molecular dynamics trajectories, enabling the segmentation of molecular conformations based on their structural similarity, typically quantified by computing the root-mean-square-distance (RMSD) between various conformations post-alignment, thereby eliminating translational and rotational discrepancies. This approach is pivotal in revealing the specific molecular conformations accessible [47, 48].

Our analysis incorporated two distinct clustering algorithms: Hierarchical Density-Based Spatial Clustering of Applications with Noise (HDBSCAN)[49] and the Amorim-Hennig method[50].

HDBSCAN, a non-parametric clustering technique, is particularly adept at analyzing disordered systems. It identifies distinct conformational clusters without necessitating a predefined number of clusters or uniform cluster density. This adaptability is crucial for explorations with undefined cluster numbers and variable densities[48, 51]

Conversely, the Amorim-Hennig method is more suitable for well-ordered systems like folded proteins. It effectively captures subtle structural changes, including alterations in secondary structural elements, and is proficient in identifying both localized and comprehensive conformational transitions[48]. The synergistic application of HDBSCAN and the Amorim-Hennig method facilitates a thorough examination of system stability and notable conformational shifts, independent of prior system-specific knowledge[48, 20, 21, 22, 23, 27, 24, 31, 33].

In this study, HDBSCAN was applied to analyze the 60s loop and the catalytic triad, while the Amorim-Hennig method was utilized for the 220s loop and the 170s loop, in line with their recognized importance in previous research. To enhance computational efficiency, our analysis was limited to the heavy atoms, omitting hydrogens, in these specified regions. Clustering was executed on the integrated 32-*µ*s trajectory from the Wake Forest University (WFU) High-Performance Computing Facility [52]. We sampled conformations every 1 ns, resulting in a total of 32,000 conformations—8,000 per system.

For visualization, thrombin structures were rendered using Tachyon in VMD 1.9.3[28]. We employed the NewCartoon representation in transparent green for representative structures, with other conformations in the same ensemble depicted as shadows. This approach visually encapsulates the extent of the conformational ensemble[53]. The median structure within each cluster, closest to the cluster’s average, was selected as the representative. Adhering to our visualization protocols[53], shadows were confined to conformations within one standard deviation of the median, effectively illustrating the variability within each cluster.

### Principal Component Analysis (PCA)

Principal Component Analysis (PCA) is an indispensable technique in the study of biological systems, particularly effective in identifying conformational variations. By reducing the dimensionality of data, PCA allows for the capturing of significant conformational changes using fewer features, thus simplifying the analysis and interpretation of complex datasets[54]. A comprehensive exposition of PCA in the context of biological systems is detailed in our recent review[47].

The PCA process in our study was executed as follows:

1. **Matrix Construction:** We formulated a matrix *A* with *T* rows and 3*N* columns, where *T* is the total number of frames in the simulation, and *N* the number of atoms in the system. Each row in *A* represents the concatenated atomic coordinates for a single frame, depicted as (*X*_1_*, Y*_1_*, Z*_1_*, X*_2_*, Y*_2_*, Z*_2_*, . . ., X_N_, Y_N_, Z_N_*).
2. **Covariance Calculation:** The covariance matrix *C* of *A* was computed using the equation:

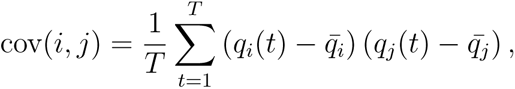

where *q* represents the coordinates *X*, *Y*, or *Z*, and *i* and *j* vary from 1 to *N* . Here, *q̄_i_* and *q̄_j_* are the time-averaged positions of *q_i_* and *q_j_*, respectively.

**3. Covariance Diagonalization:** The covariance matrix *C* was diagonalized to yield the eigenvectors *V*, forming the basis for new features. The projections *Y* = *AV* correspond to the new coordinates in this basis. Eigenvalues indicate the significance of these features, with larger values signifying greater variation in the data along those components.

In this research, PCA was implemented using the ’pca’ module in the PYEmma Python package[55], focusing on regions of interest including exosite I, the 180s loop, and the *γ* loop, and considering only the heavy atoms. Principal components were calculated based on the concatenated trajectory.

Subsequently, we generated conformational free energy landscapes for these regions. The free energy Δ*G_i_*for each bin was calculated as:

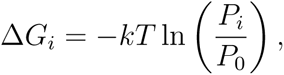

where *k* is the Boltzmann constant, *T* the temperature, *P_i_* the occupancy in the *i*-th bin, and *P*_0_ the maximum occupancy. In line with our previous work[25, 27, 20, 21, 22, 23], a binning resolution of 50 bins per dimension was employed.

### Hydrogen Bond Analysis

Hydrogen bonds play a crucial role in stabilizing the secondary and three-dimensional (3D) structures of proteins through their strong and directionally specific interactions[56, 57, 58]. These bonds are integral in maintaining protein conformations and often act as primary drivers of conformational changes[56, 57, 58].

We conducted a comprehensive hydrogen bond analysis using Python[59] and the MD-Analysis package[60, 61]. The analysis was performed on the concatenated trajectory, with hydrogen bonds identified based on a maximum heavy-atom distance of 3.2 Å and a donor-hydrogen-acceptor angle not exceeding 120°[62], in line with criteria established in our previous work[22, 23].

To improve computational efficiency, we excluded hydrogen bonds that were either infrequent or ubiquitous, as their sparse or constant presence was deemed unlikely to significantly impact our analysis. We applied conservative frequency thresholds of 2.5% for minimum and 97.5% for maximum occurrence, thus reducing approximately 80% of bonds considered non-essential. This approach decreased the computational burden fivefold. Given that the computational complexity of the logistic regression model scales linearly (*O*(*n*)) with the number of hydrogen bonds, this filtration was deemed necessary[63, 64, 22, 23].

Post-filtration, we retained a total of 509 hydrogen bonds as explanatory variables. These were represented in a structured two-dimensional matrix, with each column denoting a specific hydrogen bond and each row a distinct frame from the concatenated trajectory. Entries in this matrix were binary (1 or 0), indicating the presence or absence of a hydrogen bond in a given frame. The response variable was categorically defined to differentiate thrombin states: ’WT’ for wild-type, ’WT-TM456’ for TM456-bound wild-type, ’WE’ for the double mutant, and ’WE-TM456’ for TM456-bound double mutant. This matrix was used as the primary dataset for subsequent logistic regression analysis.

For the analysis, logistic regression was employed as the statistical model, consistent with our prior methodology[22]. This framework uses a logistic function to model the relationship between a binary response variable and one or more predictor variables[65, 66]. The logistic function maps real-valued inputs to an output range of 0 to 1, providing a probabilistic interpretation of the outcome.

The logistic regression model is mathematically represented as:

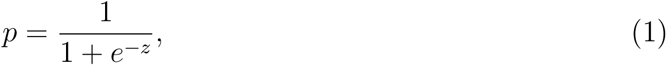

where *p* is the predicted probability of the binary response, *z* is a linear combination of predictor variables and their coefficients, and *e* is the base of the natural logarithm.

The linear combination is given by:

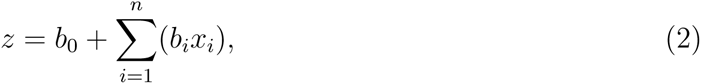

where *b*_0_ is the intercept, and *b_i_*are coefficients for each explanatory variable *x_i_*, representing the presence or absence of a hydrogen bond.

The objective of logistic regression is to optimize these coefficients to best fit the observed data, typically by minimizing the elastic net penalized negative log-likelihood. Post-training, the coefficients indicate the importance of each hydrogen bond, with larger absolute values signifying greater relevance. For a detailed exploration of this analytical method, readers are directed to our recent publications[22, 23].

## Results

### Atomic Flexibility Analysis of Thrombin: Impact of W215A/E217A Double Mutation and TM456 Binding

The atomic flexibility of thrombin, particularly under the influence of the W215A/E217A double mutation and TM456 binding, was thoroughly investigated. This analysis was centered around the Root Mean Square Fluctuations (RMSFs) of the protein’s *α*-carbons, encompassing different thrombin variants: wild-type, TM456-bound wild-type, double mutant, and TM456-bound double mutant.

Figure 1 illustrates these RMSF variations, highlighting significant changes in atomic flexibility due to the double mutation and TM456 binding. A notable increase in the flexibility of the 60s loop (comprising residues 82 to 94) was observed in both the double mutant and TM456-bound systems, indicating potential conformational dynamics.

**Figure 1:**
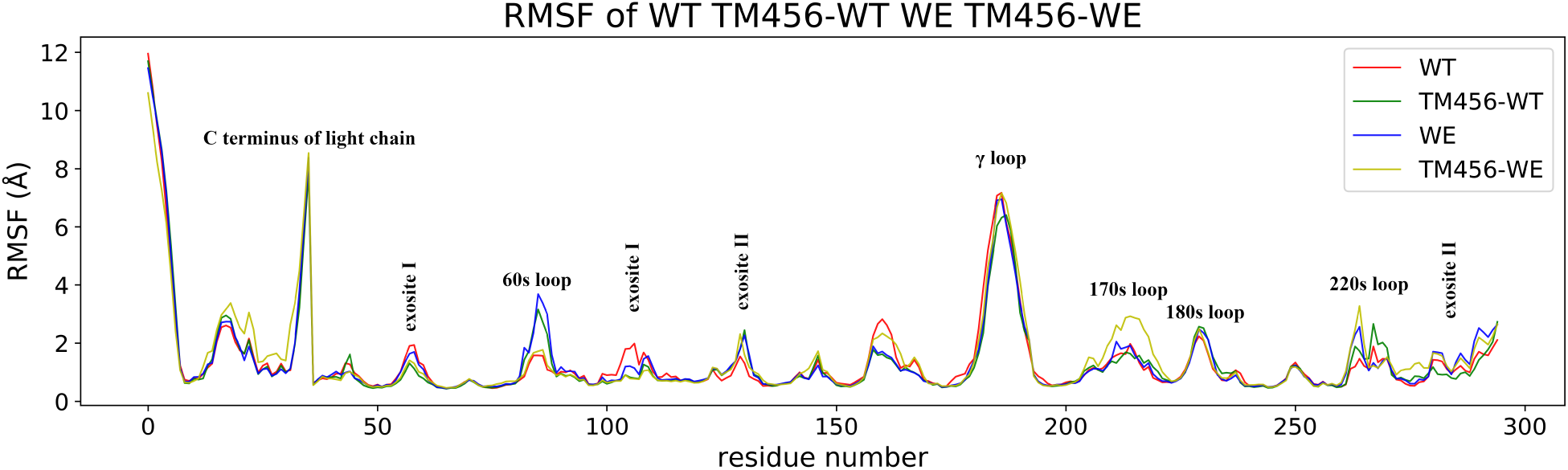
Root-mean-square fluctuations (RMSF) for the *α*-carbons of the wild-type thrombin, TM456-bound wild-type thrombin, double mutant thrombin, and TM456-bound double mutant thrombin.

**Figure 2:**
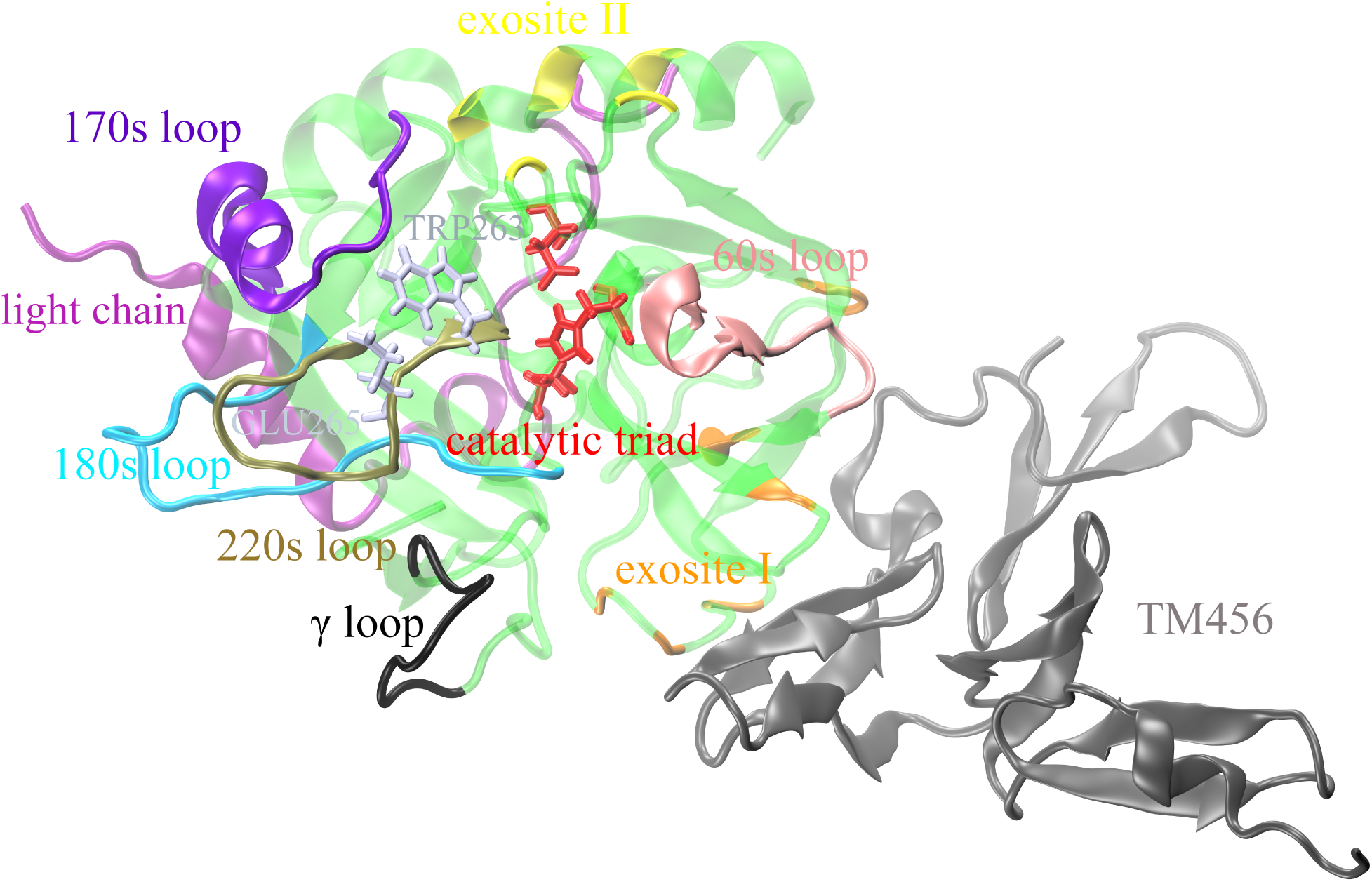
Illustration of the Thrombin Structure in Complex with TM456. Key functional regions are highlighted, including the 60s loop (pink), 170s loop (violet), 180s loop (cyan), 220s loop (tan), and the *γ* loop (black), with exosite I and II in orange and yellow, respectively. The catalytic triad side chains are in red, and mutations at TRP263 and GLU265 are shown in iceblue.

Additionally, RMSF analysis reveals that exosite I (comprising residues 57, 98, 104, 106, 109, 142, 143) experiences significant stabilization upon TM456 binding in both WT and WE systems. Interestingly, a similar stabilization trend is observed in the WE system without TM456 binding, suggesting possible allosteric effects due to the distant location of the mutation from exosite I.

The 170s loop (residues 204 to 219) demonstrates increased atomic flexibility in the WE system with TM456 binding, indicative of system-specific effects. Likewise, the 220s loop (residues 262 to 274) exhibits varying flexibility levels between WT and WE systems, with TM456 binding generally increasing flexibility in this region.

Figure 7 provides a detailed visual representation of the thrombin structure in complex with TM456. To further investigate thrombin’s conformational dynamics, a series of analytical techniques were employed, including clustering, correlation, hydrogen bond, and principal component analyses. The results from HDBSCAN are presented in Figures 3-4, while the Amorim-Hennig clustering outcomes are shown in Figures 5-6. Principal component analysis, essential for identifying primary data variations, is depicted in Figures 7-9, and hydrogen bond analysis is detailed in Figure 10. These analyses collectively offer deep insights into the conformational behavior of thrombin under the impact of the W215A/E217A double mutation and TM456 binding.

### Complex Conformational Patterns in Thrombin: Exploration via HDBSCAN and Amorim-Hennig Methods

HDBSCAN clustering revealed distinctive conformational shifts in the 60s loop, influenced by the double mutation W215A/E217A and TM456 binding, as illustrated in Figure 3(a). The systems with WE and TM456-WT exhibited a notable conformation characterized by a 180-degree rightward twist in the 60s loop (cluster 1, Figure 3(b)), absent in the WT and TM456-WE systems. This pattern indicates an independent influence of the double mutation and TM456 binding on the 60s loop, consistent with RMSF analysis results. In contrast, the TM456-WE system predominantly exhibits conformations in cluster 0, signifying a closed 60s loop (Figure 3(b)), with none in clusters 1 or 2, unlike the 0.3% and 3.64% observed in the WT. This suggests that TM456 binding to the double mutant restricts the 60s loop’s ability to open, potentially explaining the minor 7-fold reduction in protein C activation with TM presence[14, 5, 15, 16]. The consistency of the HDBSCAN results for the 60s loop with our prior study[23] (93.7% and 89.99% conformations of WT and TM456-WT in cluster 0 previously, versus 93.06% and 88.81% currently) underscores the method’s reliability for this specific system.

**Figure 3:**
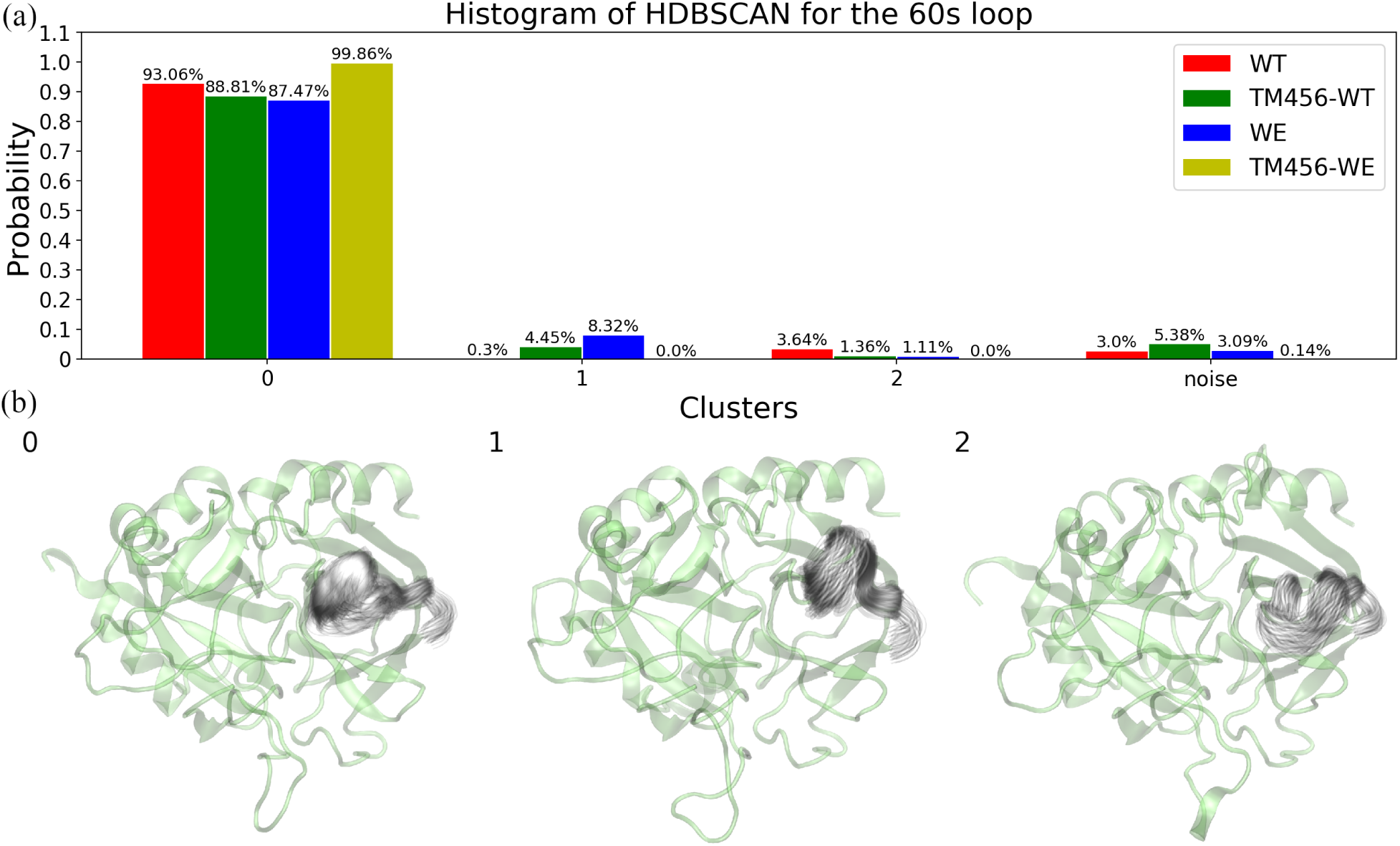
HDBSCAN Clustering of the Heavy Atoms in the 60s Loop across Different Thrombin States. (a) Distribution of clusters for the wild-type thrombin, TM456-bound wild-type thrombin, double mutant thrombin, and TM456-bound double mutant thrombin. (b) Representative structures corresponding to each cluster. The median structure of thrombin is represented via transparent green while the variance of the 60s loop in each cluster is represented by shadows according to the visual statistics[53].

The HDBSCAN analysis of the catalytic triad revealed significant conformational shifts in TM456-WT, WE, and TM456-WE systems (Figure 4). Cluster 1 contained 22.2%, 24.6%, and 16.2% of the conformations in these systems, respectively, a stark contrast to the 3.1% in wild-type thrombin (Figure 4(a)). This shift potentially elucidates the double mutant’s over 19,000-fold decrease in fibrinogen cleavage capability versus a 7-fold reduction in protein C activation[14, 5, 15, 16]. The catalytic triad exhibits two primary conformations: one in cluster 0 (Figure 4(b)), linked to fibrinogen cleavage, and another in cluster 1 (Figure 4(b)), associated with protein C activation. This distinction is supported by mutagenesis studies showing a moderate correlation (r=0.79) between thrombin’s activities towards protein C and fibrinogen [18], suggesting conformational differences for these functions. Kinetic measurements of chromogenic substrate hydrolysis by WE, with saturating thrombomodulin amounts, show a modest improvement in the *k*_cat_*/K_m_* ratio, in contrast to the 57,000-fold increase with protein C as a substrate [15, 17, 5], indicating different conformations of the catalytic triad for these substrates.

**Figure 4:**
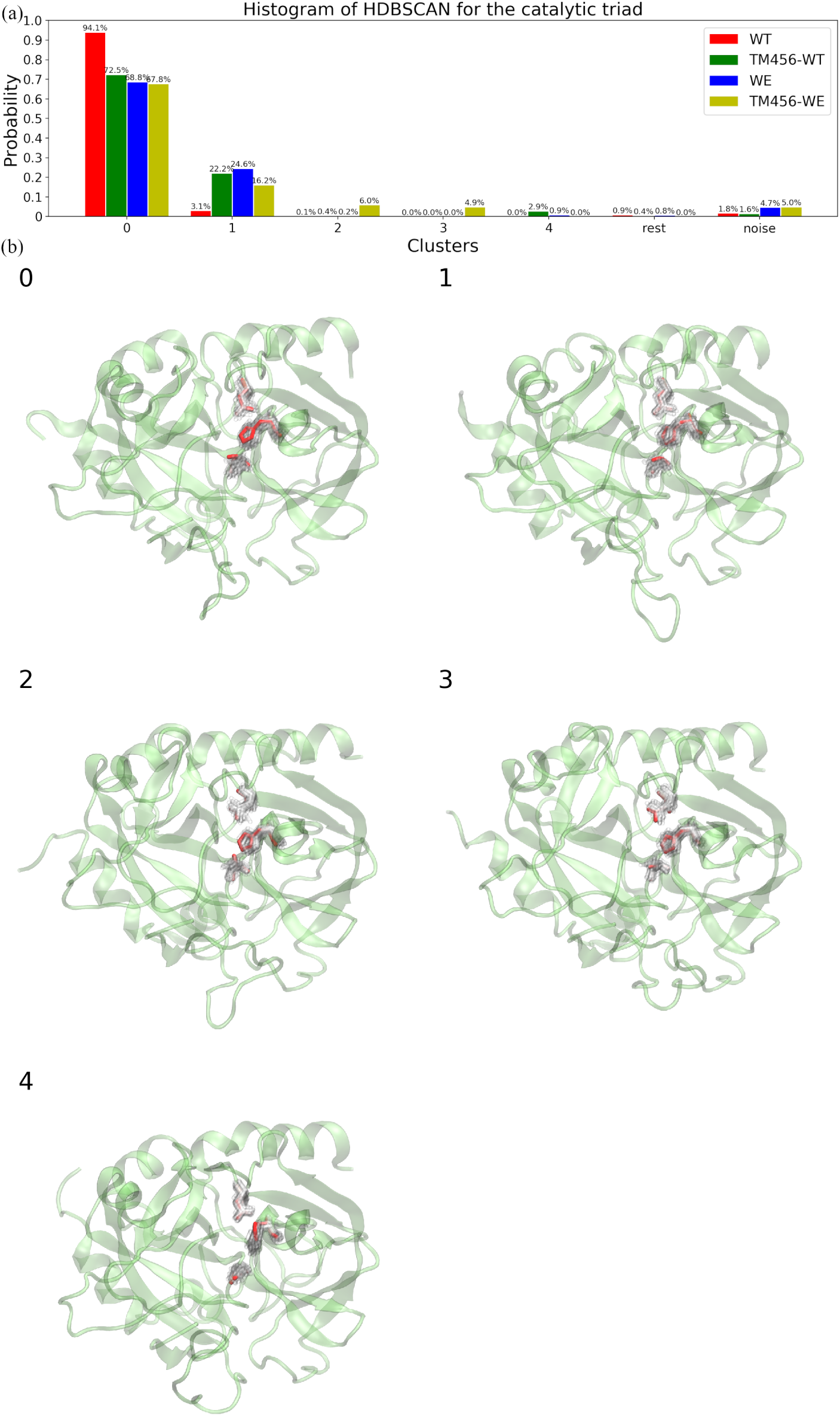
HDBSCAN Clustering of the Heavy Atoms in the Catalytic Triad Across Various Thrombin States. (a) Distribution of clusters for wild-type thrombin, TM456-bound wild-type thrombin, double mutant thrombin, and TM456-bound double mutant thrombin. (b) Representative structures corresponding to each cluster are illustrated. The median structure of thrombin is depicted in transparent green, and the variance within the catalytic triad for each cluster is represented by shadows, in accordance with the visual statistics method[53].

The 7-fold decrease in protein C activation in the TM456-WE double mutant could be related to 6% and 4.9% of conformations in clusters 2 and 3, respectively, a pattern not observed in the TM456-WT system. The HDBSCAN results for the catalytic triad, consistent with previous studies[23], support the hypothesis that the catalytic triad’s conformation in cluster 1 is crucial for protein C activation.

Amorim-Hennig clustering revealed shifts in the 220s loop in both WE and TM456-WE systems (Figure 5). The conformational distribution in clusters 0 and 1 was similar for both systems, totaling 79.72% and 82.72%, and 20.27% and 17.27%, respectively (Figure 5(a)). This indicates that TM456 binding to the double mutant does not significantly alter the 220s loop dynamics, aligning with RMSF analysis. HDBSCAN analysis of the 220s loop (Figure S1) corroborated these findings and previously reported shifts in the TM456-WT system[23], emphasizing the reliability of both clustering methods.

**Figure 5:**
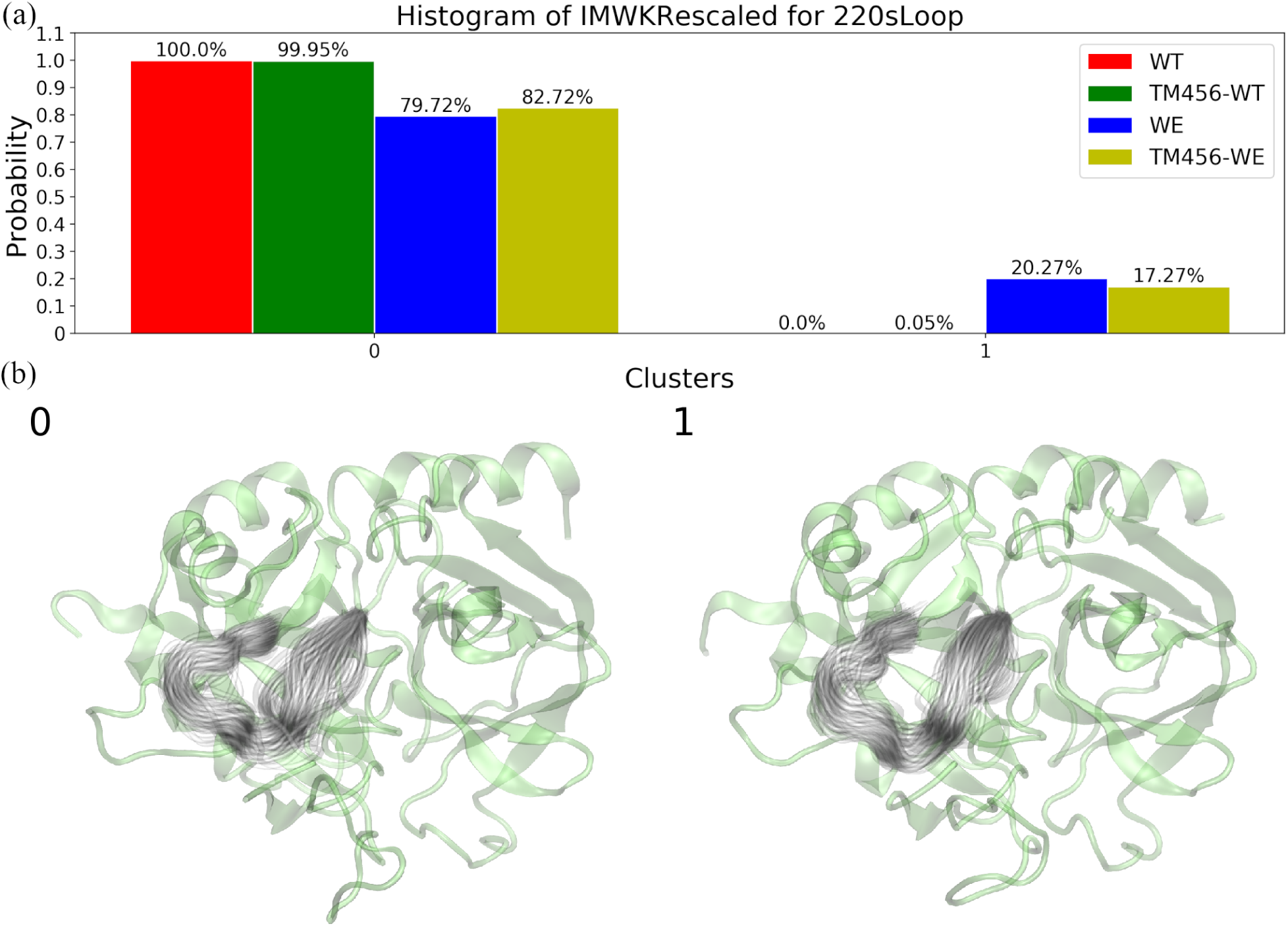
Amorim-Hennig Clustering of the Heavy Atoms in the 220s Loop across Different Thrombin States. (a) Distribution of clusters for the wild-type thrombin, TM456-bound wild-type thrombin, double mutant thrombin, and TM456-bound double mutant thrombin. (b) Representative structures corresponding to each cluster.

**Figure 6:**
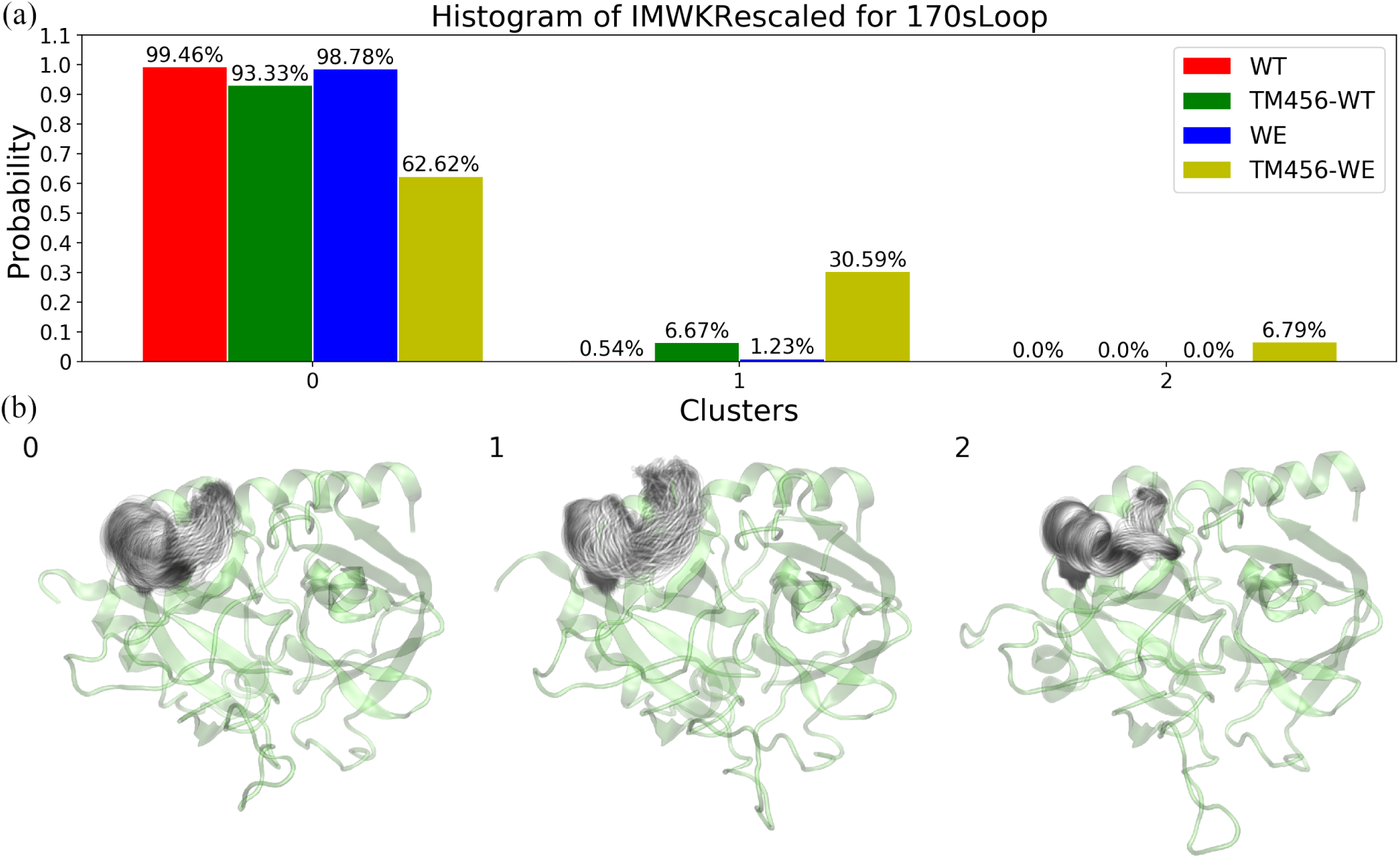
Amorim-Hennig Clustering of the Heavy Atoms in the 170s Loop across Different Thrombin States. (a) Distribution of clusters for the wild-type thrombin, TM456-bound wild-type thrombin, double mutant thrombin, and TM456-bound double mutant thrombin. (b) Representative structures corresponding to each cluster. The median structure of thrombin is represented via transparent green, while the variance of the 170s loop in each cluster is represented by shadows according to the visual statistics[53].

The Amorim-Hennig clustering method also highlighted distinct shifts in the 170s loop in both TM456-WT and TM456-WE systems, in agreement with our previous findings[23]. Specifically, 6.67% and 30.59% of the conformations in the TM456-WT and TM456-WE systems were in cluster 1, respectively. Moreover, the TM456-WE system showed 6.79% of conformations in cluster 2, a unique configuration for this loop. These shifts may explain the increased atomic flexibility observed in the 170s loop in the TM456-WE system and contribute to understanding the observed 7-fold reduction in protein C activation[14, 5, 15, 16].

### Principal Component Analysis of Exosite I, 180s, and ***γ*** Loops: Insights into TM456 Binding and Double Mutation Effects

Principal Component Analysis (PCA) reveals a notable stabilization in exosite I following TM456 binding, as evidenced by a diminished free energy well (Figure 7). This result aligns with the established role of exosite I as TM456’s binding site. In contrast, wild-type thrombin exhibits an additional free energy well, indicating a conformation potentially favorable for diverse biomolecular interactions. Interestingly, the double mutant thrombin presents this additional well in a diminished form compared to the wild-type, suggesting that mutations in the 220s loop might have allosteric effects on exosite I. This observation underscores the complex structural interplay within thrombin. Figures S2(a) and S3 visually detail these conformational differences in exosite I.

**Figure 7:**
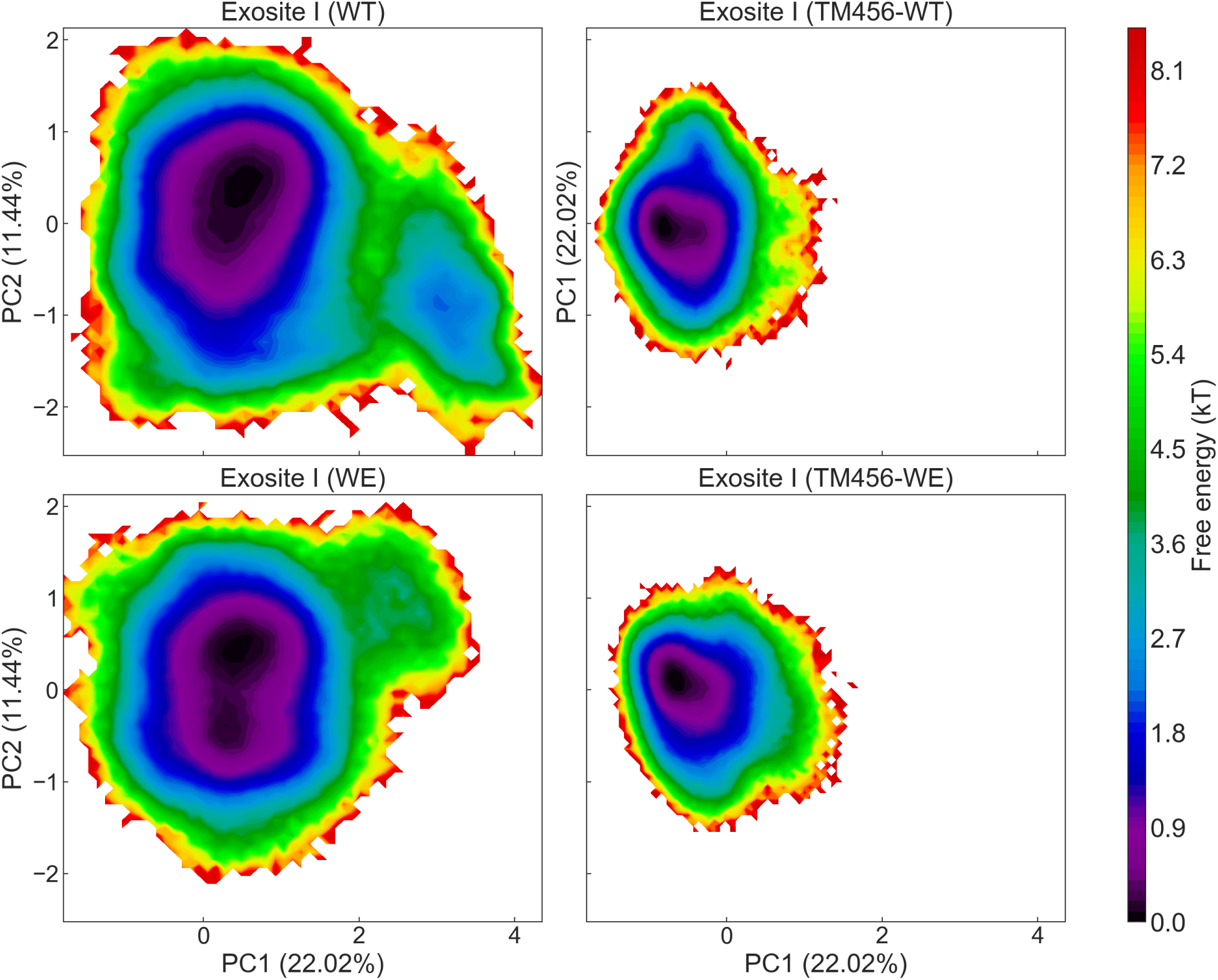
Conformational Free Energy Surfaces of Exosite I for Wild-Type Thrombin, TM456-Bound Wild-Type Thrombin, Double Mutant Thrombin, and TM456-Bound Double Mutant Thrombin.

The 180s loop is typically the most stable within thrombin’s structure. However, the binding of TM456 to wild-type thrombin induces a new free energy well for this loop (Figure 8). In the double mutant, the configuration of the free energy well undergoes a significant change, merging three wells into two. This change likely relates to the spatial proximity of the 180s and 220s loops, where the double mutation resides. These alterations highlight the complex interconnectivity of thrombin’s structural elements. Figures S2(b) and S4 illustrate the varied conformations of the 180s loop within these wells.

**Figure 8:**
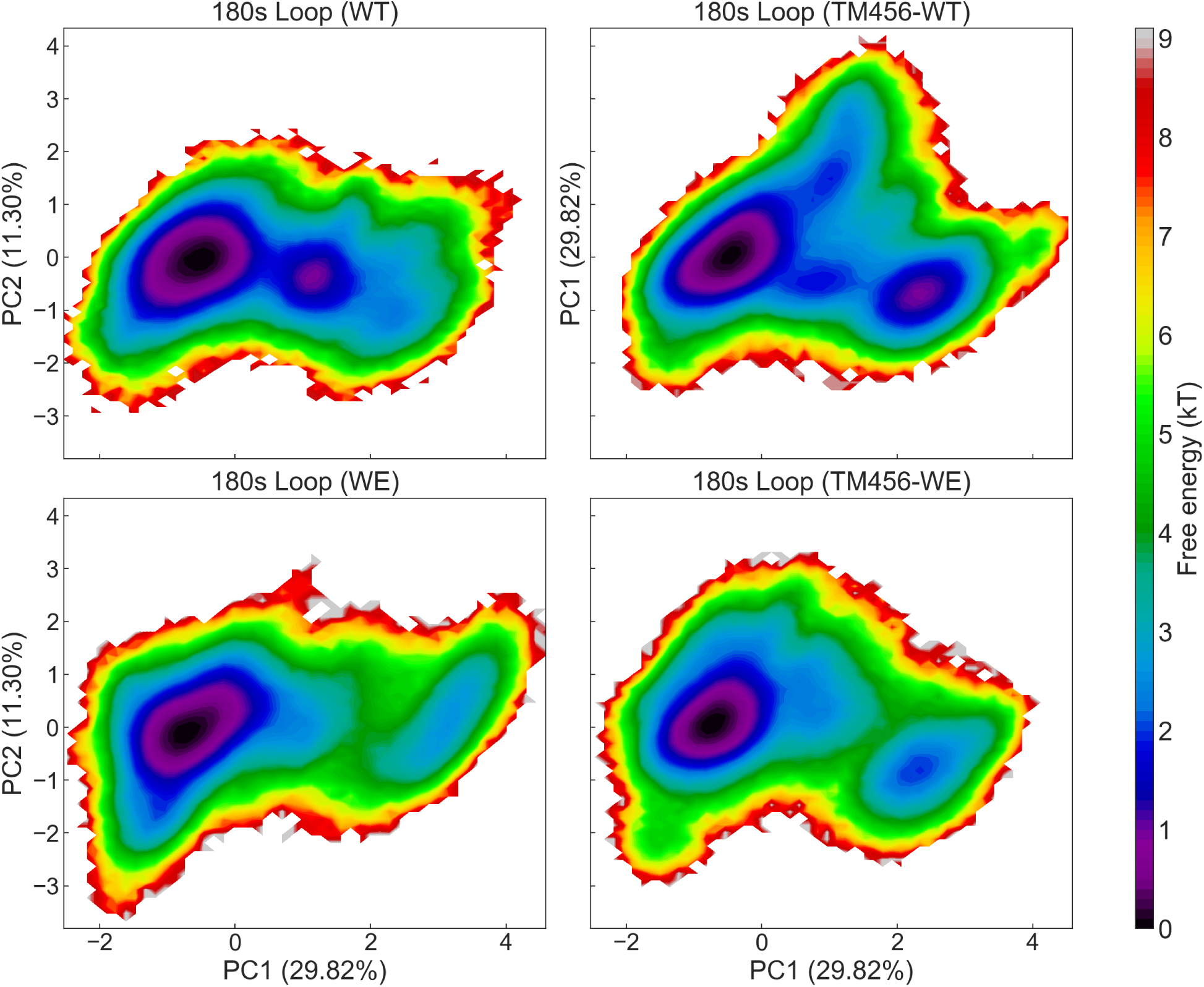
Conformational Free Energy Surfaces of the 180s Loop for Wild-Type Thrombin, TM456-Bound Wild-Type Thrombin, Double Mutant Thrombin, and TM456-Bound Double Mutant Thrombin.

Moreover, PCA delineates the distinct effects on the *γ* loop’s free energy well from TM456 binding and the double mutation (Figure 9). TM456 binding compresses the well’s contour while increasing conformational dispersion within it. Conversely, the double mutation alone shows minimal impact on the well, suggesting only slight alterations in the *γ* loop’s conformations. However, TM456 binding to the double mutant centralizes the *γ* loop’s conformations within a narrower well. This interaction, particularly evident in the TM456-WE systems, may offer crucial insights into the mechanistic basis for the 7-fold reduction in protein C activation in these systems[14, 5, 15, 16]. The diverse conformations of the *γ* loop, distributed across seven different wells, are comprehensively depicted in Figures S2(c) and S5.

**Figure 9:**
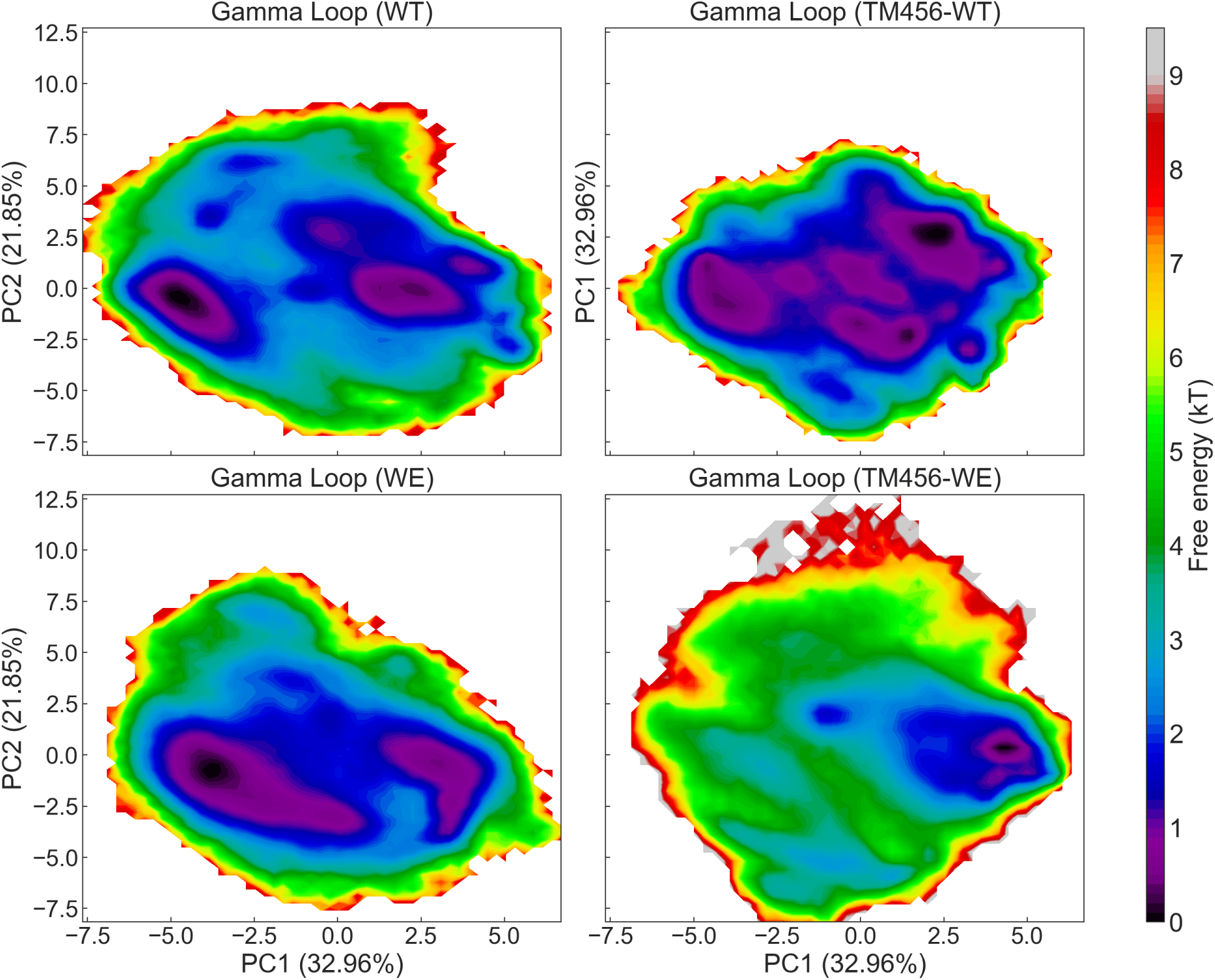
Conformational Free Energy Surfaces of the *γ* Loop for Wild-Type Thrombin, TM456-Bound Wild-Type Thrombin, Double Mutant Thrombin, and TM456-Bound Double Mutant Thrombin.

### Logistic Regression-Based Exploration of Allosteric Hydrogen Bonds in TM456-Bound and Double Mutant Thrombin

Logistic regression models were employed to differentiate between wild-type thrombin, TM456-bound wild-type thrombin, double mutant thrombin, and TM456-bound double mutant thrombin. These models exhibited prediction accuracies of 97.09%, 98.03%, and 98.71%, respectively. Remarkably, even when the models were constrained to only the top 13, 12, and 11 hydrogen bonds, they achieved accuracies of 87.26%, 86.38%, and 89.61%, respectively. This suggests that over 90% of the original prediction accuracy is maintained, underscoring the pivotal role of these specific hydrogen bonds as discriminative features. Figure 10 graphically represents these hydrogen bonds, and Table 1 enumerates their coefficients in the logistic regression models.

**Figure 10:**
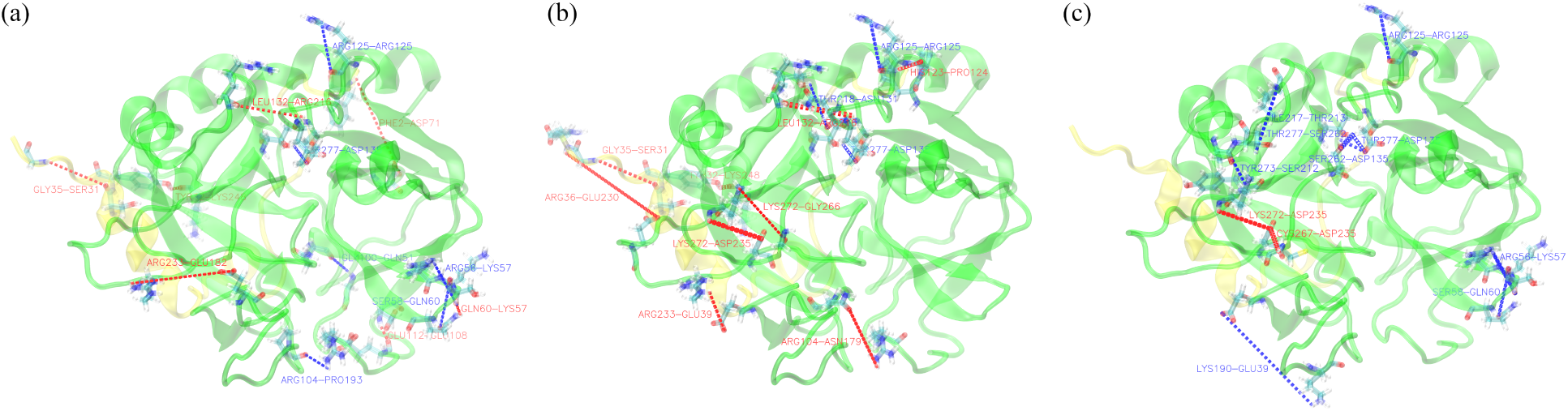
Visualization of critical hydrogen bonds identified through logistic regression models, specific to (a) TM456-bound wild-type thrombin, (b) double mutant thrombin, and (c) TM456-bound double mutant thrombin. The light and heavy chains of thrombin are portrayed using NewCartoon representation and are rendered in transparent yellow and green, respectively. Relevant residues are displayed using the Licorice representation. Hydrogen bonds are marked by dashed lines: blue lines indicate negative beta values, signifying hydrogen bonds that form upon TM456 binding or the introduction of double mutations, while red lines signify positive beta values, denoting hydrogen bonds that break following TM456 binding or double mutations. The line thickness is proportional to the absolute value of the beta values. Labels for each hydrogen bond are positioned directly adjacent to the corresponding dashed lines.

**Table 1:**
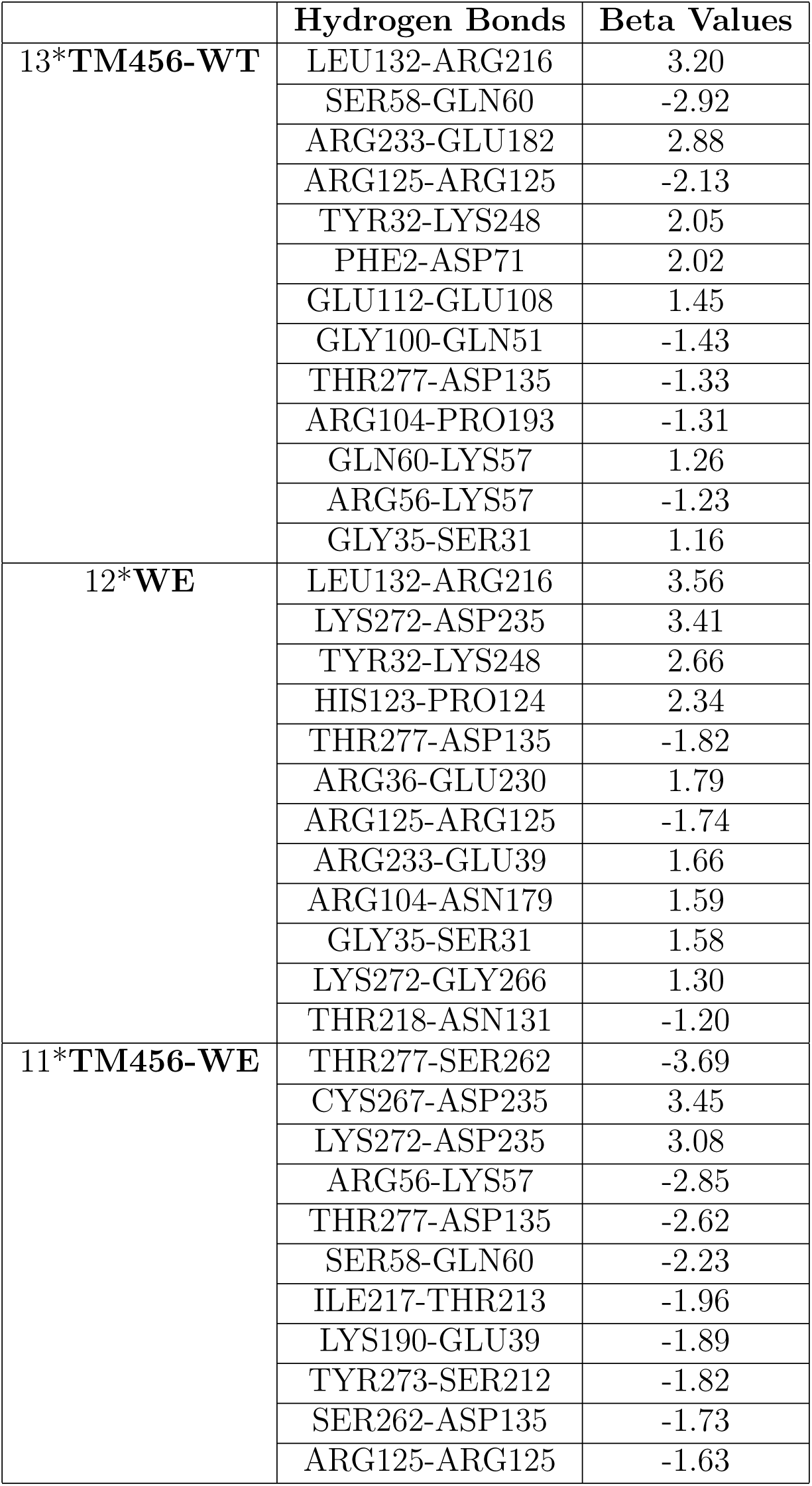
Logistic Regression Beta Values for Hydrogen Bonds Across Thrombin Variants: WT-TM456, WE, and WE-TM456.

|  | Hydrogen Bonds | Beta Values |
| --- | --- | --- |
| **13\*TM456-WT** | LEU132-ARG216 | 3.20 |
|  | SER58-GLN60 | -2.92 |
|  | ARG233-GLU182 | 2.88 |
|  | ARG125-ARG125 | -2.13 |
|  | TYR32-LYS248 | 2.05 |
|  | PHE2-ASP71 | 2.02 |
|  | GLU112-GLU108 | 1.45 |
|  | GLY100-GLN51 | -1.43 |
|  | THR277-ASP135 | -1.33 |
|  | ARG104-PRO193 | -1.31 |
|  | GLN60-LYS57 | 1.26 |
|  | ARG56-LYS57 | -1.23 |
|  | GLY35-SER31 | 1.16 |
| **12\*WE** | LEU132-ARG216 | 3.56 |
|  | LYS272-ASP235 | 3.41 |
|  | TYR32-LYS248 | 2.66 |
|  | HIS123-PRO124 | 2.34 |
|  | THR277-ASP135 | -1.82 |
|  | ARG36-GLU230 | 1.79 |
|  | ARG125-ARG125 | -1.74 |
|  | ARG233-GLU39 | 1.66 |
|  | ARG104-ASN179 | 1.59 |
|  | GLY35-SER31 | 1.58 |
|  | LYS272-GLY266 | 1.30 |
|  | THR218-ASN131 | -1.20 |
| **11\*TM456-WE** | THR277-SER262 | -3.69 |
|  | CYS267-ASP235 | 3.45 |
|  | LYS272-ASP235 | 3.08 |
|  | ARG56-LYS57 | -2.85 |
|  | THR277-ASP135 | -2.62 |
|  | SER58-GLN60 | -2.23 |
|  | ILE217-THR213 | -1.96 |
|  | LYS190-GLU39 | -1.89 |
|  | TYR273-SER212 | -1.82 |
|  | SER262-ASP135 | -1.73 |
|  | ARG125-ARG125 | -1.63 |

Despite the disparate spatial locations of TM456 binding and double mutations, both TM456-bound wild-type thrombin and double mutant thrombin exhibit similar effects on the catalytic triad, as illustrated in Figures 10(a) and 10(b). This congruence is partly attributed to shared allosteric pathways, particularly through the hydrogen bonds `LEU132-ARG216` and `THR277-ASP135`. Additional hydrogen bonds, notably `ARG125-ARG125` and `GLY35-SER31`, are significant for both forms, further reinforcing their similar biochemical impacts. Unique hydrogen bonds such as `LYS272-ASP235` and `LYS272-GLY266`, emerging due to double mutations around the 220s loop, and `ARG56-LYS57`, `SER58-GLN60`, `GLN60-LYS57`, and `GLU112-GLU108`, resulting from TM456 binding near exosite I, serve to differentiate these forms.

In contrast to expectations, the essential hydrogen bonds in TM456-bound double mutant thrombin (Figures 10(c)) do not represent a simple superposition of those identified as significant in the other forms. The allosteric pathway modulating the catalytic triad in this form involves a unique series of hydrogen bonds—`THR277-SER262`, `SER262-ASP135`, and `THR277-ASP135`—distinct from the `LEU132-ARG216` and `THR277-ASP135` bonds observed in the other forms. The key bond `THR277-ASP135` remains constant, linked to the catalytic triad residue `ASP135`. This is supported by HDBSCAN clustering analysis of the catalytic triad, revealing a close alignment of conformations in cluster 1 for TM456-bound double mutant thrombin with those in the other forms, while clusters 2 and 3 likely represent unique allosteric pathways.

Additionally, specific hydrogen bonds like `ILE217-THR213` and `TYR273-SER212` in TM456-bound double mutant thrombin, correlated with structural changes in the 170s loop (residues 204 to 219), are identified. Hydrogen bonds such as `ARG56-LYS57`, `SER58-GLN60`, `LYS272-ASP235`, and `CYS267-ASP235` appear induced by TM456 binding and double mutations, resembling those in the other thrombin forms.

Each thrombin form—TM456-bound wild-type, double mutant, and TM456-bound double mutant—features a unique hydrogen bond (`LYS190-GLU39`, `ARG104-PRO193`, and `ARG104-ASN179`, respectively) related to allosteric changes in the *γ* loop. These findings are consistent with PCA analysis of the *γ* loop’s free energy well. Notably, `ARG125-ARG125` is the only hydrogen bond consistently significant across all forms, aside from `THR277-ASP135`, suggesting a potential influence on the structural dynamics of the catalytic triad and identifying `ARG125` as a potential therapeutic target.

## Conclusions

In this study, we utilized a comprehensive suite of advanced computational methods, including Root Mean Square Fluctuation (RMSF), clustering algorithms (HDBSCAN and Amorim-Hennig), Principal Component Analysis (PCA), and logistic regression models. Our focus was on dissecting the intricate structural dynamics and allosteric regulatory mechanisms of various thrombin forms: wild-type, TM456-bound wild-type, WE, and TM456-bound WE. The central aim was to comprehend how the WE double mutant, despite having a compromised catalytic pocket, retains its proficiency in activating protein C when coupled with Thrombomodulin (TM).

Our analysis revealed the likely existence of two distinct conformations within the catalytic triad of thrombin. One conformation is specialized for fibrinogen cleavage, while the other facilitates the activation of protein C. This bifurcation in functionality was initially suggested by HDBSCAN clustering and further corroborated by an in-depth analysis of hydrogen bonding patterns, aligning with multiple experimental studies[18, 15, 17, 5]. Notably, the WE double mutations appear to allosterically shift the catalytic triad from its fibrinogen-cleaving state in the wild-type to a state conducive to protein C activation. Consequently, the WE double mutant demonstrates a significant 19,000-fold reduction in fibrinogen cleavage capabilities[14, 5, 15, 16]. However, in the presence of TM, this mutant exhibits only a minor 7-fold decrease in protein C activation efficiency[14, 5, 15, 16], indicative of the catalytic triad adopting a conformation favorable for protein C activation. Additional distinctive conformations observed in TM456-bound WE, such as those in the catalytic triad located in clusters 2 and 3, the 170s loop conformations in clusters 1 and 2, the unique free energy landscape of the *γ* loop, and the conserved 60s loop, collectively contribute to the observed reduction in protein C activation efficiency.

Furthermore, our findings highlight a synergistic interplay between the WE mutations and TM456 binding in altering thrombin’s structural dynamics. This synergy is evidenced by the observation that the critical hydrogen bonds in TM456-bound WE, as identified through logistic regression analysis, do not simply represent a combination of those significant in TM456-bound wild-type and WE thrombin. The binding of TM456 to the WE mutant uniquely alters the free energy landscape of the *γ* loop, diverging from the individual influences exerted by either the WE mutations or TM456 binding alone. This provides additional evidence of their synergistic effects. Moreover, TM456-bound WE displays unique conformations, including specialized ones in the catalytic triad found in clusters 2 and 3, and distinct 170s loop conformations categorized within clusters 1 and 2. These observations collectively support the hypothesis that the WE mutations and TM456 binding synergistically impact thrombin’s structural and functional characteristics.

Additionally, our study identified both unique and shared key hydrogen bonds across the different thrombin variants. A significant finding was the consistent importance of the hydrogen bond involving `ARG125` across all thrombin variants, suggesting its potential as a therapeutic target.

In summary, this study provides a detailed understanding of thrombin’s allosteric regulations influenced by WE mutations and TM456 binding. By employing sophisticated computational methodologies, we have elucidated the conformational states and highlighted the synergistic effects that enable the WE double mutant to maintain functional capabilities despite structural constraints. These insights not only advance our knowledge of thrombin’s biological mechanisms but also suggest new avenues for targeted therapeutic strategies, particularly highlighting `ARG125` as a promising therapeutic target.

## Data and Software Availability

All associated code utilized in this study is readily accessible at our GitHub repository: https://github.com/salsburygroup/Thrombin_TM_W215A_E217A. Due to their substantial size, the raw trajectory files are not directly available online. However, they can be obtained upon request.

## Acknowledgements

The authors wish to acknowledge the support of the Wake Forest Baptist Comprehensive Cancer Center Crystallography & Computational Biosciences Shared Resource, supported by the National Cancer Institute’s Cancer Center Support Grant award number P30CA012197. The content is solely the responsibility of the authors and does not necessarily represent the official views of the National Cancer Institute. Portions of the computations were performed on the Wake Forest University DEAC Cluster, a centrally managed resource with support provided in part by Wake Forest University[52]. FRS would also like to thank the Scott Family for the Scott Family Fellowship. DW would like to thank the Center for Molecular Signaling at Wake Forest University for a fellowship.

## References

[1] Earl W Davie and John D Kulman. “An overview of the structure and function of thrombin”. In: Seminars in thrombosis and hemostasis. Vol. 32. S 1. Copyright© 2006 by Thieme Medical Publishers, Inc., 333 Seventh Avenue, New . . . 2006, pp. 003–015.

[2] Enrico Di Cera. “Thrombin”. In: Mol. Aspects Med. 29.4 (2008), pp. 203–254.

[3] Caroline J Reddel, Chuen Wen Tan, and Vivien M Chen. “Thrombin Generation and Cancer: Contributors and Consequences”. en. In: Cancers 11.1 (Jan. 2019), p. 100.

[4] Allison S Remiker and Joseph S Palumbo. “Mechanisms coupling thrombin to metastasis and tumorigenesis”. en. In: Thromb. Res. 164 Suppl 1 (Apr. 2018), S29–S33.

[5] Prafull S Gandhi, Michael J Page, Zhiwei Chen, Leslie Bush-Pelc, and Enrico Di Cera. “Mechanism of the anticoagulant activity of thrombin mutant W215A/E217A”. In: Journal of Biological Chemistry 284.36 (2009), pp. 24098–24105.

[6] Andras Gruber, Angelene M Cantwell, Enrico Di Cera, and Stephen R Hanson. “The thrombin mutant W215A/E217A shows safe and potent anticoagulant and antithrombotic effects in vivo”. In: Journal of Biological Chemistry 277.31 (2002), pp. 27581– 27584.

[7] Andras Gruber, JA Fernandez, L Bush, U Marzec, JH Griffin, SR Hanson, and E Di Cera. “Limited generation of activated protein C during infusion of the protein C activator thrombin analog W215A/E217A in primates”. In: Journal of Thrombosis and Haemostasis 4.2 (2006), pp. 392–397.

[8] Michelle A Berny, Tara C White, Erik I Tucker, Leslie A Bush-Pelc, Enrico Di Cera, Andŕas Gruber, and Owen JT McCarty. “Thrombin mutant W215A/E217A acts as a platelet GPIb antagonist”. In: Arteriosclerosis, thrombosis, and vascular biology 28.2 (2008), pp. 329–334.

[9] Cristina P Vicente, Hartmut Weiler, Enrico Di Cera, and Douglas M Tollefsen. “Thrombomodulin is required for the antithrombotic activity of thrombin mutant W215A/E217A in a mouse model of arterial thrombosis”. In: Thrombosis research 130.4 (2012), pp. 646–648.

[10] Erik I Tucker, Norah G Verbout, Brandon D Markway, Michael Wallisch, Christina U Lorentz, Monica T Hinds, Joseph J Shatzel, Leslie A Pelc, David C Wood, Owen JT McCarty, et al. “The protein C activator AB002 rapidly interrupts thrombus development in baboons”. In: *Blood*, The Journal of the American Society of Hematology 135.9 (2020), pp. 689–699.

[11] Michelle A Berny-Lang, Sawan Hurst, Erik I Tucker, Leslie A Pelc, Ruikang K Wang, Patricia D Hurn, Enrico Di Cera, Owen JT McCarty, and Andŕas Gruber. “Thrombin mutant W215A/E217A treatment improves neurological outcome and reduces cerebral infarct size in a mouse model of ischemic stroke”. In: Stroke 42.6 (2011), pp. 1736–1741.

[12] CS Gibbs, SE Coutre, M Tsiang, WX Li, AK Jain, KE Dunn, VS Law, CT Mao, SY Matsumura, SY Mejza, et al. “Conversion of thrombin into an anticoagulant by protein engineering”. In: Nature 378.6555 (1995), pp. 413–416.

[13] Daniele Arosio, Youhna M Ayala, and Enrico Di Cera. “Mutation of W215 compromises thrombin cleavage of fibrinogen, but not of PAR-1 or protein C”. In: Biochemistry 39.27 (2000), pp. 8095–8101.

[14] Kenichi A Tanaka, Andras Gruber, Fania Szlam, Leslie A Bush, Stephen R Hanson, and Enrico Di Cera. “Interaction between thrombin mutant W215A/E217A and direct thrombin inhibitor”. In: Blood coagulation & fibrinolysis: an international journal in haemostasis and thrombosis 19.5 (2008), p. 465.

[15] Angelene M Cantwell and Enrico Di Cera. “Rational design of a potent anticoagulant thrombin”. In: Journal of Biological Chemistry 275.51 (2000), pp. 39827–39830.

[16] Riley B Peacock, Taylor McGrann, Sofia Zaragoza, and Elizabeth A Komives. “How thrombomodulin enables W215A/E217A thrombin to cleave protein C but not fibrinogen”. In: Biochemistry 61.2 (2022), pp. 77–84.

[17] Agustin O Pineda, Zhi-Wei Chen, Sonia Caccia, Angelene M Cantwell, Savvas N Savvides, Gabriel Waksman, F Scott Mathews, and Enrico Di Cera. “The anticoagulant thrombin mutant W215A/E217A has a collapsed primary specificity pocket”. In: Journal of Biological Chemistry 279.38 (2004), pp. 39824–39828.

[18] Enrico Di Cera. “Thrombin as an anticoagulant”. In: Progress in molecular biology and translational science 99 (2011), pp. 145–184.

[19] Pablo Fuentes-Prior, Yoriko Iwanaga, Robert Huber, Rene Pagila, Galina Rumennik, Marian Seto, John Morser, David R Light, and Wolfram Bode. “Structural basis for the anticoagulant activity of the thrombin–thrombomodulin complex”. In: Nature 404.6777 (2000), pp. 518–525.

[20] Dizhou Wu, Jiajie Xiao, and Freddie R Salsbury Jr. “Light Chain Mutation Effects on the Dynamics of Thrombin”. In: Journal of Chemical Information and Modeling 61.2 (2021), pp. 950–965.

[21] Dizhou Wu and Freddie R Salsbury. “Simulations suggest double sodium binding induces unexpected conformational changes in thrombin”. In: Journal of Molecular Modeling 28.5 (2022), pp. 1–16.

[22] Dizhou Wu and Freddie R. Salsbury. “Unraveling the Role of Hydrogen Bonds in Thrombin via Two Machine Learning Methods”. In: Journal of Chemical Information and Modeling (June 2023). doi: 10.1021/acs.jcim.3c00153. url: https://doi.org/10.1021/acs.jcim.3c00153.

[23] Dizhou Wu, Athul Prem, Jiajie Xiao, and Freddie R Salsbury. “Thrombin-A Molecular Dynamics Perspective.” In: Mini Reviews in Medicinal Chemistry (2023).

[24] Jiajie Xiao, Ryan L Melvin, and Freddie R Salsbury. “Mechanistic Insights Into Thrombin’s Switch Between “Slow” and “Fast” Forms”. In: Phys. Chem. Chem. Phys. 19.36 (2017), pp. 24522–24533.

[25] Jiajie Xiao and Freddie R Salsbury. “Molecular Dynamics Simulations of Aptamer-Binding Reveal Generalized Allostery in Thrombin”. In: J. Biomol. Struct. Dyn. 35.15 (2017), pp. 3354–3369.

[26] Jiajie Xiao and Freddie R Salsbury. “Na+-Binding Modes Involved in Thrombin’s Allosteric Response as Revealed by Molecular Dynamics Simulations, Correlation Networks and Markov Modeling”. In: Phys. Chem. Chem. Phys. 21.8 (2019), pp. 4320–4330.

27. Jiajie Xiao, Ryan L Melvin, and Freddie R Salsbury Jr. “Probing Light Chain Mutation Effects on Thrombin via Molecular Dynamics Simulations and Machine Learning”. In: J. Biomol. Struct. Dyn. 37.4 (2019), pp. 982–999.

[28] William Humphrey, Andrew Dalke, and Klaus Schulten. “Vmd: Visual Molecular Dynamics”. In: J. Mol. Graphics 14.1 (1996), pp. 33–38.

[29] Matt J Harvey, Giovanni Giupponi, and G De Fabritiis. “ACEMD: Accelerating Biomolecular Dynamics in the Microsecond Time Scale”. In: J. Chem. Theory Comput. 5.6 (2009), pp. 1632–1639.

[30] Ryan L Melvin, William H Gmeiner, and Freddie R Salsbury Jr. “All-Atom molecular dynamics reveals mechanism of zinc complexation with therapeutic F10”. In: The journal of physical chemistry B 120.39 (2016), pp. 10269–10279.

[31] Ryan L Melvin, William G Thompson, Ryan C Godwin, William H Gmeiner, and Freddie R Salsbury Jr. “Muts*α*’s Multi-Domain Allosteric Response to Three Dna Damage Types Revealed by Machine Learning”. In: Front. Phys.(Lausanne) 5 (2017), p. 10.

[32] Ryan L Melvin, William H Gmeiner, and Freddie R Salsbury Jr. “All-Atom MD Predicts Magnesium-Induced Hairpin in Chemically Perturbed RNA Analog of F10 Therapeutic”. In: The Journal of Physical Chemistry B 121.33 (2017), pp. 7803–7812.

[33] Ryan L Melvin, William H Gmeiner, and Freddie R Salsbury. “All-atom MD indicates ion-dependent behavior of therapeutic DNA polymer”. In: Physical Chemistry Chemical Physics 19.33 (2017), pp. 22363–22374.

[34] Ryan Godwin, William Gmeiner, and Freddie R Salsbury Jr. “Importance of Long-Time Simulations for Rare Event Sampling in Zinc Finger Proteins”. In: J. Biomol. Struct. Dyn. 34.1 (2016), pp. 125–134.

[35] Ryan C Godwin, Ryan L Melvin, William H Gmeiner, and Freddie R Salsbury Jr. “Binding Site Configurations Probe the Structure and Dynamics of the Zinc Finger of Nemo (Nf-*κ*B Essential Modulator)”. In: Biochemistry 56.4 (2017), pp. 623–633.

[36] Ryan C Godwin, Lindsay M Macnamara, Rebecca W Alexander, and Freddie R Salsbury Jr. “Structure and Dynamics of tRNAMet Containing Core Substitutions”. In: ACS omega 3.9 (2018), pp. 10668–10678.

[37] Ryan C Godwin, William H Gmeiner, and Freddie R Salsbury Jr. “All-atom molecular dynamics comparison of disease-associated zinc fingers”. In: Journal of Biomolecular Structure and Dynamics 36.10 (2018), pp. 2581–2594.

[38] William L Jorgensen, Jayaraman Chandrasekhar, Jeffry D Madura, Roger W Impey, and Michael L Klein. “Comparison of Simple Potential Functions for Simulating Liquid Water”. In: J. Chem. Phys. 79.2 (1983), pp. 926–935.

[39] Jing Huang and Alexander D MacKerell Jr. “CHARMM36 all-atom additive protein force field: Validation based on comparison to NMR data”. In: Journal of computational chemistry 34.25 (2013), pp. 2135–2145.

[40] Herman JC Berendsen, JPM van Postma, Wilfred F van Gunsteren, ARHJ DiNola, and Jan R Haak. “Molecular Dynamics With Coupling to an External Bath”. In: J. Chem. Phys. 81.8 (1984), pp. 3684–3690.

[41] Don S Lemons and Anthony Gythiel. “Paul Langevin’s 1908 Paper “on the Theory of Brownian Motion”[“sur la théorie du mouvement brownien,” cr acad. sci.(Paris) 146, 530–533 (1908)]”. In: Am. J. Phys. 65.11 (1997), pp. 1079–1081.

[42] Tom Darden, Darrin York, and Lee Pedersen. “Particle Mesh Ewald: An N Log (N) Method for Ewald Sums in Large Systems”. In: J. Chem. Phys. 98.12 (1993), pp. 10089–10092.

[43] MJ Harvey and G De Fabritiis. “An Implementation of the Smooth Particle Mesh Ewald Method on Gpu Hardware”. In: J. Chem. Theory Comput. 5.9 (2009), pp. 2371–2377.

[44] K Anton Feenstra, Berk Hess, and Herman JC Berendsen. “Improving Efficiency of Large Time-Scale Molecular Dynamics Simulations of Hydrogen-Rich Systems”. In: J. Comput. Chem. 20.8 (1999), pp. 786–798.

[45] WF Van Gunsteren and Herman JC Berendsen. “Algorithms for Macromolecular Dynamics and Constraint Dynamics”. In: Mol. Phys. 34.5 (1977), pp. 1311–1327.

[46] Salsbury Group. AMD Python: An implementation of the Accelerated Molecular Dynamics in Python. https://github.com/salsburygroup/amd-python. Accessed: 2023-07-17. 2023.

[47] Ryan C Godwin, Ryan Melvin, and Freddie R Salsbury. “Molecular Dynamics Simulations and Computer-Aided Drug Discovery”. In: Computer-aided drug discovery. Springer, 2015, pp. 1–30.

[48] Ryan L Melvin, Ryan C Godwin, Jiajie Xiao, William G Thompson, Kenneth S Berenhaut, and Freddie R Salsbury Jr. “Uncovering Large-Scale Conformational Change in Molecular Dynamics Without Prior Knowledge”. In: J. Chem. Theory Comput. 12.12 (2016), pp. 6130–6146.

[49] Ricardo J. G. B. Campello, Davoud Moulavi, and Joerg Sander. “Density-Based Clustering Based on Hierarchical Density Estimates”. In: Advances in Knowledge Discovery and Data Mining. Springer Berlin Heidelberg, 2013, pp. 160–172. doi: 10.1007/978-3-642-37456-2_14. url: https://doi.org/10.1007/978-3-642-37456-2_14.

[50] Renato Cordeiro De Amorim and Christian Hennig. “Recovering the Number of Clusters in Data Sets With Noise Features Using Feature Rescaling Factors”. In: Inf. Sci. (N. Y.) 324 (2015), pp. 126–145.

[51] Ryan L Melvin, William H Gmeiner, and Freddie R Salsbury. “All-atom MD indicates ion-dependent behavior of therapeutic DNA polymer”. en. In: Phys. Chem. Chem. Phys. 19.33 (Aug. 2017), pp. 22363–22374.

[52] Information Systems and Wake Forest University. WFU High Performance Computing Facility. 2021. doi: 10.57682/G13Z-2362. url: https://hpc.wfu.edu.

[53] Ryan L Melvin and Freddie R Salsbury Jr. “Visualizing Ensembles in Structural Biology”. In: J. Mol. Graphics Modell. 67 (2016), pp. 44–53.

[54] Hervé Abdi and Lynne J Williams. “Principal Component Analysis”. In: Wiley Interdiscip. Rev. Comput. Stat. 2.4 (2010), pp. 433–459.

[55] Martin K Scherer, Benjamin Trendelkamp-Schroer, Fabian Paul, Guillermo Pérez-Herńandez, Moritz Hoffmann, Nuria Plattner, Christoph Wehmeyer, Jan-Hendrik Prinz, and Frank Nóe. “Pyemma 2: A Software Package for Estimation, Validation, and Analysis of Markov Models”. In: J. Chem. Theory Comput. 11.11 (2015), pp. 5525–5542.

[56] Lydia M Gregoret, Stephen D Rader, Robert J Fletterick, and Fred E Cohen. “Hydrogen bonds involving sulfur atoms in proteins”. In: Proteins: Structure, Function, and Bioinformatics 9.2 (1991), pp. 99–107.

[57] Alan R Fersht and Luis Serrano. “Principles of protein stability derived from protein engineering experiments”. In: Current opinion in structural biology 3.1 (1993), pp. 75–83.

[58] David Schell, Jerry Tsai, J Martin Scholtz, and C Nick Pace. “Hydrogen bonding increases packing density in the protein interior”. In: Proteins: Structure, Function, and Bioinformatics 63.2 (2006), pp. 278–282.

[59] Guido vanRossum. “Python reference manual”. In: Department of Computer Science [CS] R 9525 (1995).

[60] Naveen Michaud-Agrawal, Elizabeth J Denning, Thomas B Woolf, and Oliver Beckstein. “MDAnalysis: a toolkit for the analysis of molecular dynamics simulations”. In: Journal of computational chemistry 32.10 (2011), pp. 2319–2327.

[61] Richard J Gowers, Max Linke, Jonathan Barnoud, Tyler JE Reddy, Manuel N Melo, Sean L Seyler, Jan Domanski, David L Dotson, Śebastien Buchoux, Ian M Kenney, et al. “MDAnalysis: a Python package for the rapid analysis of molecular dynamics simulations”. In: Proceedings of the 15th python in science conference. Vol. 98. SciPy Austin, TX. 2016, p. 105.

[62] Thomas Steiner. “The hydrogen bond in the solid state”. In: Angewandte Chemie International Edition 41.1 (2002), pp. 48–76.

[63] Andrew I Schein and Lyle H Ungar. “Active learning for logistic regression: an evaluation”. In: Machine Learning 68.3 (2007), pp. 235–265.

[64] Habiba Muhammad Sani, Ci Lei, and Daniel Neagu. “Computational complexity analysis of decision tree algorithms”. In: International Conference on Innovative Techniques and Applications of Artificial Intelligence. Springer. 2018, pp. 191–197.

[65] Michael P LaValley. “Logistic regression”. In: Circulation 117.18 (2008), pp. 2395–2399.

[66] Garetsh James, Daniela Witten, Trevor Hastie, and Robert Tibshirani. An introduction to statistical learning. Vol. 112. Springer, 2013.

