## Supplemental Info for "Unraveling the Impact of W215A/E217A Mutations on Thrombin’s Dynamics and Thrombomodulin Binding through Molecular Dynamics Simulations"

### Supplementary figures

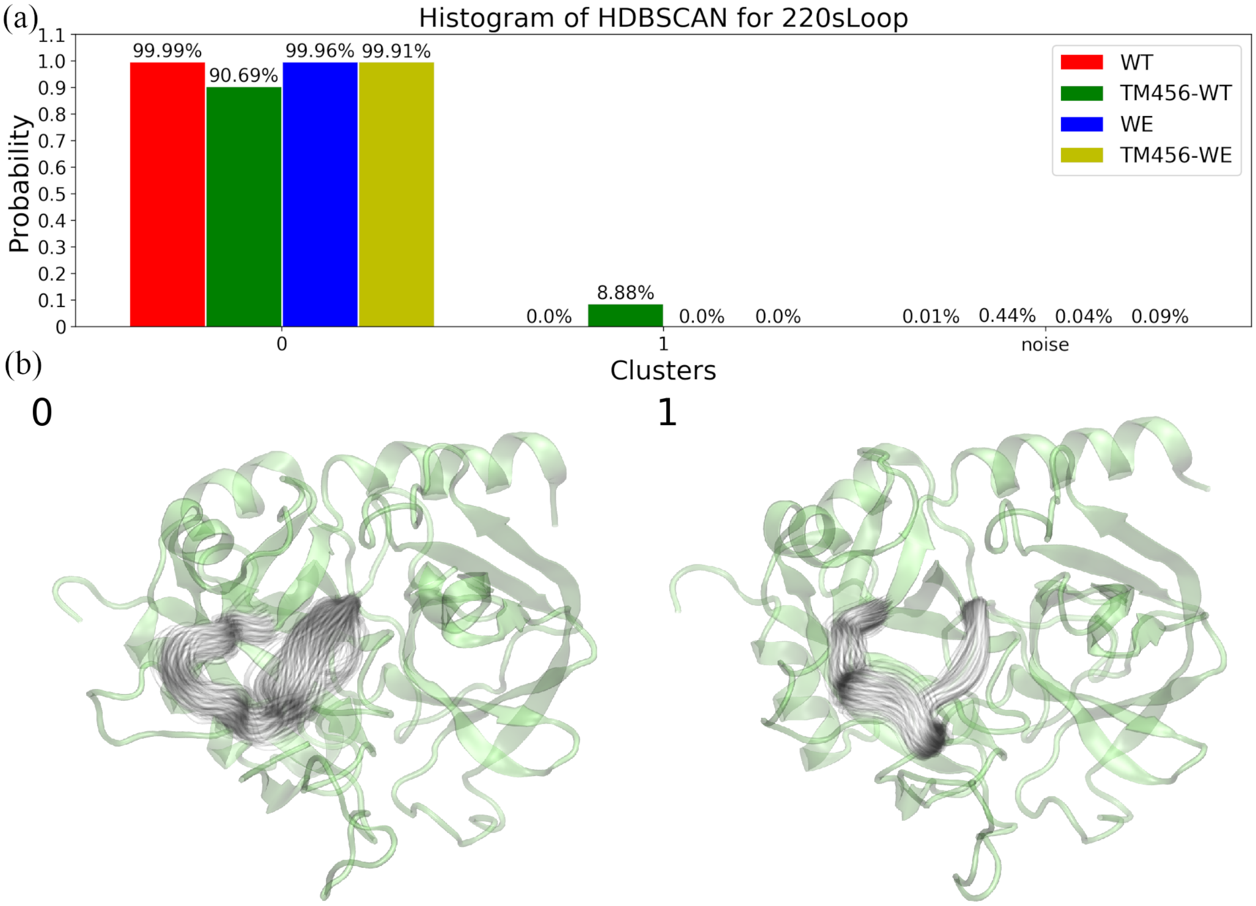


Figure S1: HDBSCAN Clustering of the Heavy Atoms in the 220s Loop across Different Thrombin States. (a) Distribution of clusters for the wild-type, TM456-bound wild-type, WE, and TM456-bound WE. (b) Representative structures corresponding to each cluster.

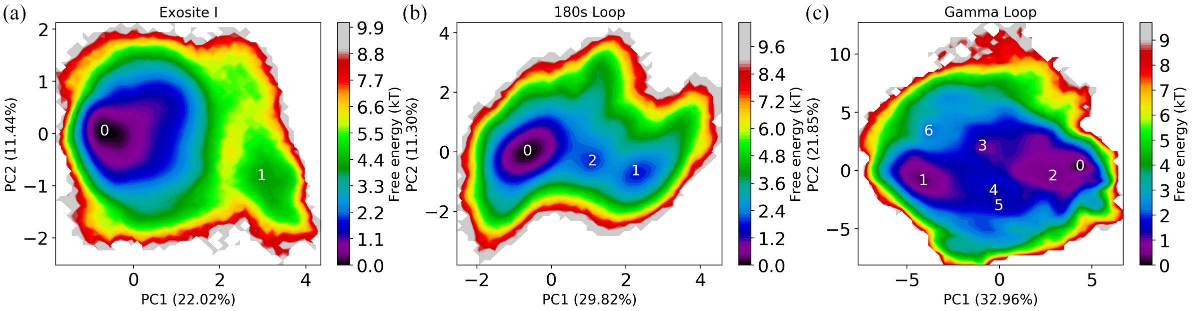


Figure S2: Conformational Free Energy Surfaces of (a) Exosite I, (b) 180s Loop, and (c) Gamma Loop for the concatenated trajectory of wild-type, TM456-bound wild-type, WE, and TM456-bound WE.

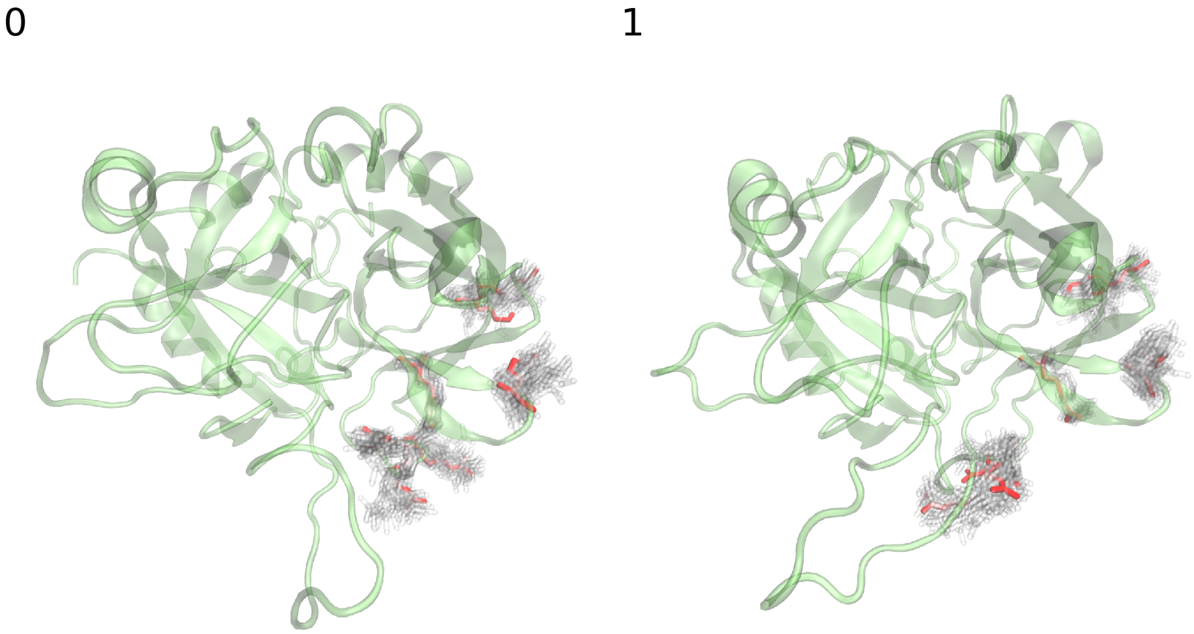


Figure S3: Representative Structures for Exosite I in Wells 0 and 1: Presented in Transparent Green using NewCartoon Representation. Red Licorice Depictions Indicate Sidechains within Exosite I. Accompanying Gray Shadows Illustrate Variance in Exosite I.

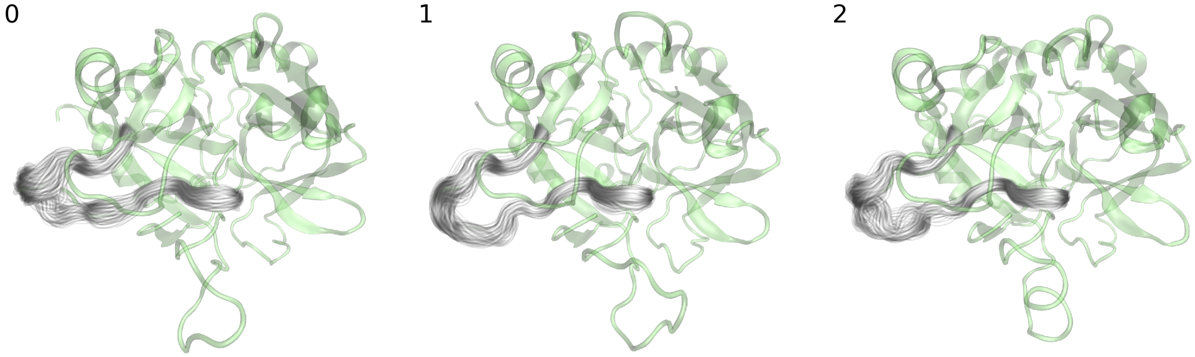


Figure S4: Representative Structures for 180s Loop in Wells 0 to 2: Presented in Transparent Green using NewCartoon Representation. Accompanying Gray Shadows Illustrate Variance in 180s Loop.

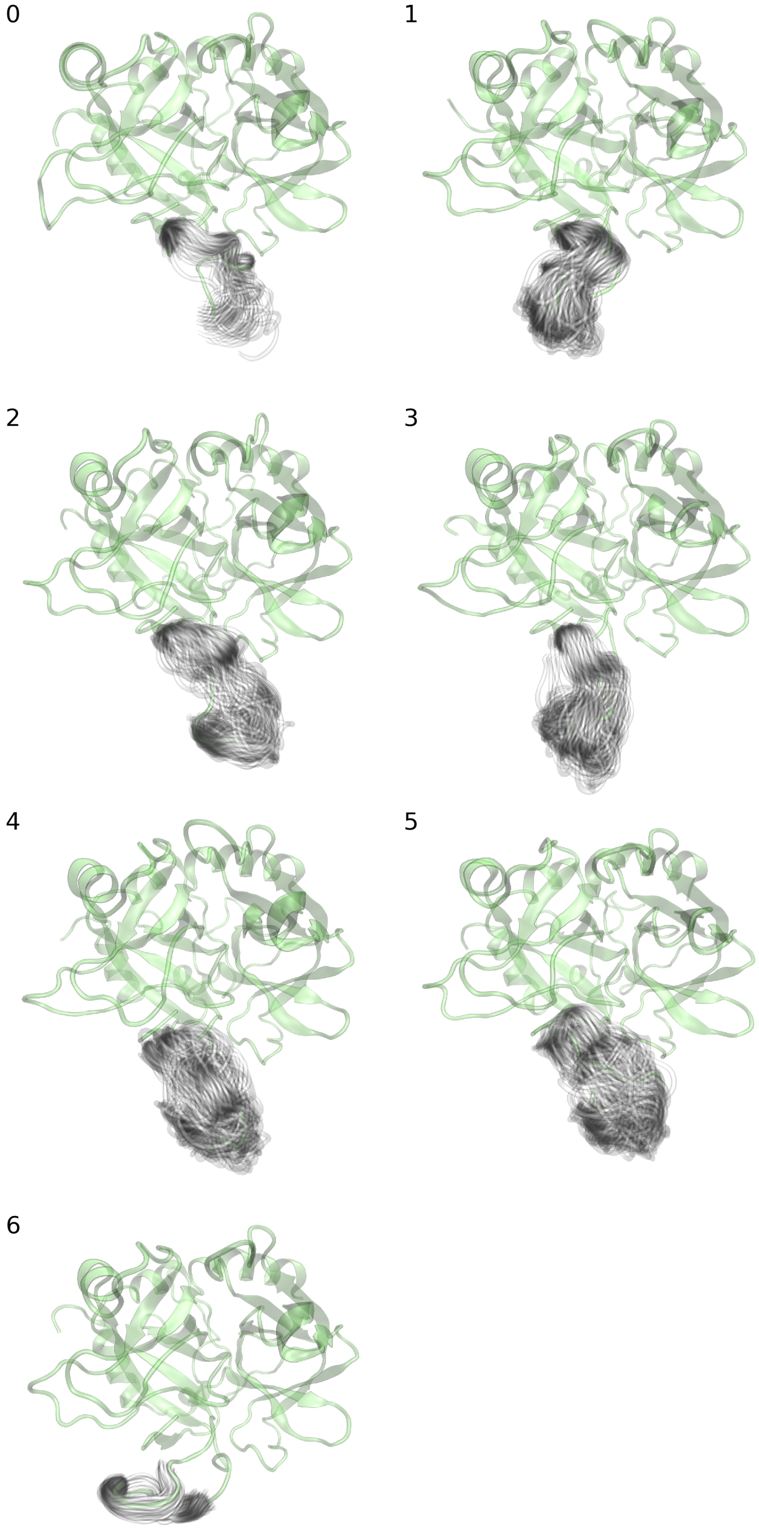


Figure S5: Representative Structures for Gamma Loop in Wells 0 to 6: Presented in Transparent Green using NewCartoon Representation. Accompanying Gray Shadows Illustrate Variance in Gamma Loop.
